# Genomic evidence for recurrent genetic admixture during domestication mediterranean olive trees (*Olea europaea*)

**DOI:** 10.1101/2020.03.28.013227

**Authors:** Irene Julca, Marina Marcet-Houben, Fernando Cruz, Jèssica Gómez-Garrido, Brandon S. Gaut, Concepción M. Díez, Ivo G. Gut, Tyler S. Alioto, Pablo Vargas, Toni Gabaldón

## Abstract

**Background:** The olive tree (*Olea europaea* L. subsp. *europaea*, Oleaceae) has been the most economic perennial crop for Mediterranean countries since its domestication around 6,000 years ago. Two taxonomic varieties are currently recognized: cultivated (var. *europaea*) and wild (var. *sylvestris*) trees. To shed light into the recent evolution and domestication of the olive tree, we sequenced the genomes of twelve individuals: ten var. *europaea*, one var. *sylvestris*, and one outgroup taxon (subsp. *cuspidata*). All of them were analysed together with an improved assembly of var. *europaea* reference genome and the available assembly of var. *sylvestris*.

**Results:** Our analyses show that cultivated olives exhibit slightly lower levels of overall genetic diversity than wild forms, and that this can be partially explained by the occurrence of a mild population bottleneck 5000-7000 years ago during the primary domestication period. We also provide the first phylogenetic analysis of genome-wide sequences, which supports a continuous process of domestication of the olive tree. This, together with population structure and introgression analyses highlights genetic admixture with wild populations across the Mediterranean Basin in the course of domestication.

**Conclusions:** Altogether, our results suggest that a primary domestication area in the eastern Mediterranean basin was followed by numerous secondary events across most countries of southern Europe and northern Africa, often involving genetic admixture with genetically rich wild populations, particularly from the western Mediterranean Basin. Based on selection tests and a search for selective sweeps, we found that genes associated with stress response and developmental processes were positively selected in cultivars. However, we did not find evidence that genes involved in fruit size or oil content were under positive selection.

## Background

The Mediterranean olive tree (*Olea europaea* L. subsp. *europaea*, Oleaceae) is one of the earliest cultivated fruit trees of the Mediterranean Basin (MB). Current classifications recognize two taxonomic varieties of *O. europaea:* var. *sylvestris* (hereafter *sylvestris*, also named oleaster) for wild populations and var. *europaea* (hereafter *europaea*) for cultivated forms (1,2). Both varieties are predominantly out-crossing, and have long lifespans, including a long juvenile phase that can last up to 15 years in natural conditions. The natural distribution of the Mediterranean olive encompasses all countries of the MB, although a few wild populations have also been found in northern areas with low occurrence of frost (3). Cultivars have historically been planted nearby wild populations since ancient times, where they exchange pollen that have resulted in effective crop production and historical hybridization (4).

There is a large body of evidence pointing to the eastern MB as the cradle of the first domestication event of olives. According to archaeological, palaeobotanical, and genetic studies, the crop was domesticated from eastern wild progenitors around 6,000 years ago (5–8). Once superior individuals were selected, clonal propagation made their multiplication and the fixation of valuable agronomic traits possible. Clonal propagation also facilitated the spread of cultivars from the eastern to the western MB via classical civilizations such as Phoenicians, Greeks, or Romans. After six millennia, olive domestication has resulted in a vast number of cultivars of uncertain pedigree that are often geographically restricted to local areas (9).

It remains unclear whether olive cultivars derived from a single initial domestication event in the Levant, followed by secondary diversification (8,10,11), or whether cultivated lineages are the result of more than a single, independent primary domestication event (7,12–15). Previous studies based on plastid and nuclear markers have suggested controversial but not necessarily incompatible domestication scenarios. The genealogical reconstruction of plastid lineages have yielded unresolved phylogenies at the base of domesticated lineages due to low plastid diversity (8,16,17). For instance, 90% of olive cultivars across the MB share the same “eastern like” plastid haplotype (17). This general result is congruent with archaeological data (5), which suggests a major domestication event in the Levant, possibly followed by recurrent admixture events with local wild olives that would have contributed to the crop diversity. However, nuclear markers showed a more complex pattern. Olive cultivars clustered together into three different gene pools, with a rough geographical correspondence to the eastern, central and western MB (12,18–21). The relationship between these groups also showed interesting features (12): the western group (southern Spain and Portugal) retained the fingerprint of a genetic bottleneck, and surprisingly, was closely related to cultivated accessions from the Levant (12). The central MB group, which also included the cultivars from eastern Spain (Catalonia, Valencia and Balearic Islands), showed signals of recent and extensive admixture with local wild populations and relatively high plastid diversity compared to the other groups of cultivars. These differential patterns, along with the fact that many cultivars from the central MB retain wild-like phenotypic characteristics, opened the controversial question of a possible minor domestication center for olives in the central MB (10,14).

Approximate Bayesian Computing (ABC) models were applied to infer the demographic history of olives and were consistent with a primary domestication event in the East (8,12). These models also highlighted the paramount role of admixture to account for the diversity of the crop. This feature was particularly predominant in cultivars from the central MB, where ~20% of the genetic diversity of olives may have been acquired via introgression with local wild populations (12). So far, genetic and archeological sources of evidence have agreed with the existence of a major center of domestication for olives in the Levant, but have been insufficient to prove or discard the existence of secondary centers of domestication elsewhere in the MB. To gain novel insights into this open question and into the most recent evolution of the cultivated olive tree, we sequenced twelve accessions, including ten cultivar representatives (‘Arbequina’, ‘Beladi’, ‘Picual’, ‘Sorani’, ‘Chemlal de Kabilye’, ‘Megaritiki, ‘Lechin de Sevilla’, ‘Lechin de Granada’, ‘Frantoio’, and ‘Koroneiki’), one wild individual of var. *sylvestris*, and one individual from *O. europaea* subsp. *cuspidata* to be used as a distant outgroup in our analysis (Table 1). The ten cultivars were carefully selected to represent: (i) the main cultivar diversity from the most important areas of Mediterranean olive cultivation. For instance, ‘Arbequina’ is the most international cultivar due to its adaptation to high density planting designs, ‘Picual’ covers more than 1.5 millions of hectares in southern Spain, ‘Frantoio’ and ‘Koroneiki’ are the primary cultivars in Italy and Greece, respectively, and ‘Beladi’ and ‘Sorani’ are widely used in the Levant; (ii) the three genetic pools identified in the MB (‘Picual’ and ‘Lechin de Granada’ – West; ‘Arbequina’, ‘Frantoio’, ‘Koroneiki’ and ‘Megaritiki’ – Central; ‘Beladi’ and ‘Sorani’ – East); and (iii) the main plastid lineages found in the cultivated olive (‘Lechin de Sevilla’ and ‘Megaritiki’ – E2.3 and E2.2, respectively; ‘Chemlal de Kabilye’ – E3.2; the rest of cultivars – E1.1). For *sylvestris*, previous phylogenetic results and field experience led us to choose a wild individual from an isolated area in order to avoid potential feral or highly introgressed trees. Hence, we sampled a tree from a location by the coast near the Cantabrian mountains (northern Spain) where the olive tree has not been historically cultivated and that is 200 km distant from current plantations. These populations had previously been screened using fingerprinting techniques (3) and Sanger sequencing (17).

**Table 1.**
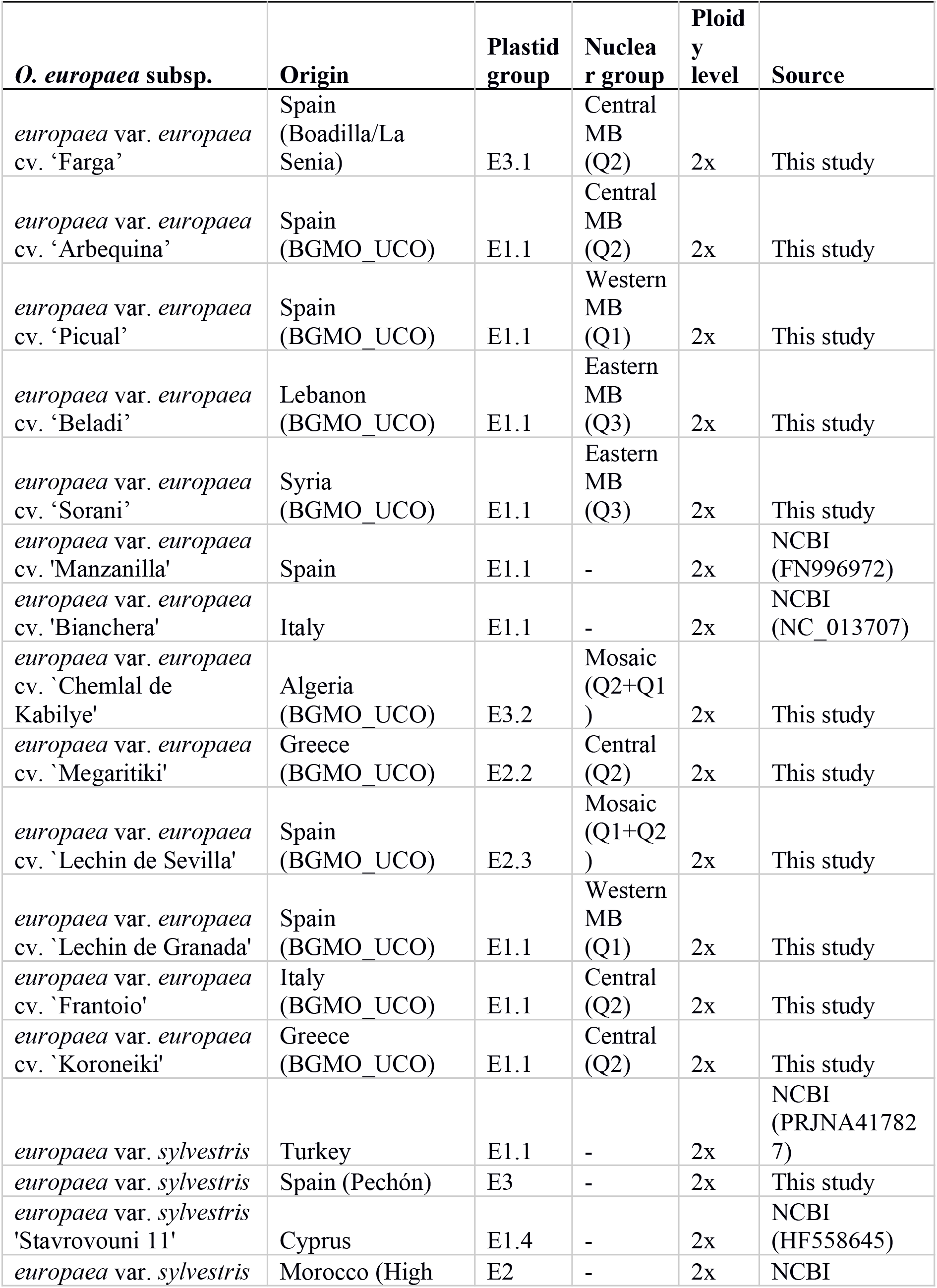
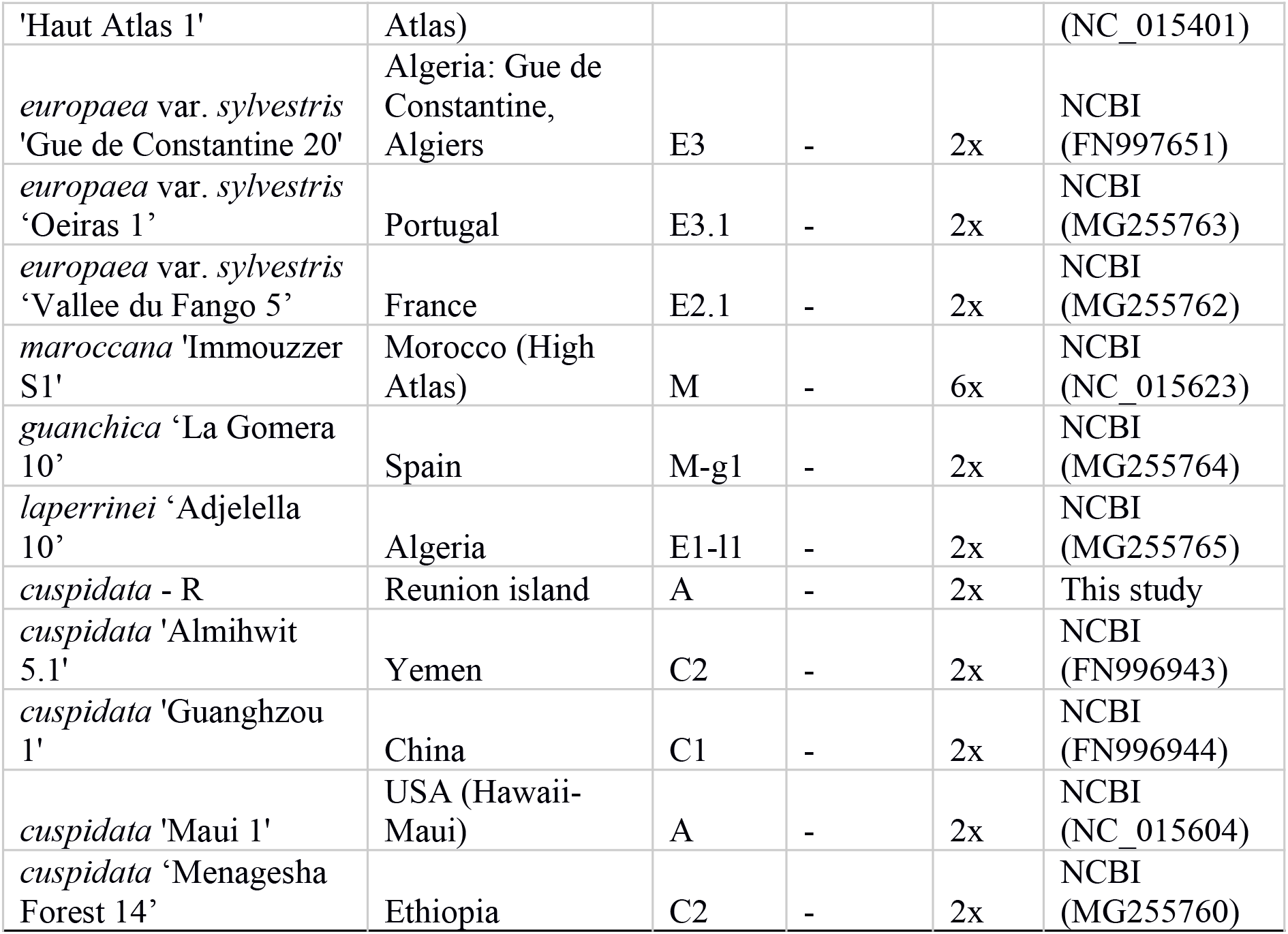
*O. europaea* genomes used in the analysis. The columns show the sample origin, plastid group (16), nuclear group (12), ploidy level, and the source of the data.

The twelve newly obtained sequences complement the two available genomes for the species: the cultivar Farga from eastern Spain, for which we provide here an improved assembly by anchoring it to a genetic map, and an oleaster (var. *sylvestris*) from Turkey (22,23). The analysis of these genomes is meant to shed light on the recent evolution and domestication of the olive tree. In particular, we addressed the question whether genome wide variation data can disentangle scenarios of one versus multiple centers of domestication. Additionally, we were interested in finding out whether genetic introgression between wild and cultivated trees have historically played a role in the domestication process, to assess the presence of potential domestication bottlenecks and to identify genes and genomic regions under selection. Finally, sequencing of nuclear genomes allowed testing earlier suggestions of close relationship between cultivars of distant locations such as southern Spain and the Levant (12).

## Results

### New assembly version of the reference olive (cultivar Farga) genome

We improved the available *O. europaea* var. *europaea* (cultivar Farga) genome assembly (Oe6 version) (22) by anchoring it to chromosomes using a publicly available genetic map (24) and removing 201 scaffolds which likely represent contaminating sequences (See materials and methods). In the final assembly (Oe9), 520.5 Mbp (39.5%) of Oe6 sequence was anchored to 23 linkage groups, of which 288 Mbp (21.8%) were oriented. This anchored assembly (Oe9) has a lower N50 as compared to the recently published assembly for *O. europaea* var. *sylvestris* (23), but it is much less fragmented, displaying a ~5 fold reduction in the number of scaffolds (9,751 vs 41,261 scaffolds, Additional file 1: Fig. S1 and Additional file 2: Table S1). The comparison between both assemblies (Additional file 1: Fig. S1b) shows a high level of conserved synteny and confirms the ancient polyploidization event in *O. europaea* described earlier (25) as many regions appear duplicated between the two genomes.

Additionally, we improved the genome annotation in Oe9 genome, by extended automated functional annotation and by manual curation of some of the genes. Based on this, the Oe9 assembly has 4,911 more annotated genes than the var. *sylvestris* genome. A comparison of gene sets using BLASTN (26) with identity >80% and e-value<1e-5 cutoffs, shows that 5,245 Oe9 annotated genes do not have a match in *sylvestris*, and, conversely, 2,620 genes of *sylvestris* do not have a match in *europaea*. Distinct genome annotation methods could partly explain these differences (22,23). To have an annotation-independent measure of differences between both assemblies, we mapped raw reads of both genome projects to the alternative assembly and assessed coverage of the putative unique genes (see material and methods). We first filtered out the genes that have at least 50% of their length with a read coverage higher than 20, which resulted in 2,115 and 280 unique genes for *europaea* and *sylvestris*, respectively. Even when lowering the coverage threshold to 5, *europaea* still had more unique genes (1,756) than *sylvestris* (102). Of these, we discarded 131 Oe9 specific genes as possible contaminations as their first BLAST hit fell outside plants. Thus, some of the genes uniquely found in the Oe9 assembly may represent true differences in terms of gene content. Interestingly, some unique genes found in *europaea* have functions associated with stress response, such as HIPPs (heavy metal-associated isoprenylated plant proteins) (27), LEA (Late Embryogenesis Abundant) (28), and salicylic acid-binding (29). Other genes are associated with growth and development. This is the case of RALF (Rapid ALkalinization Factor), which has been shown to arrest root growth and development in tomato and *Arabidopsis* (30), and caffeoyl shikimate esterase (CSE), which is an enzyme central to the lignin biosynthetic pathway (31). It is well known that lignin biosynthesis contributes to plant growth, tissue and organ development, and response to a variety of biotic and abiotic stresses (32). Some other Oe9 unique genes were associated with seed dormancy and sugar signaling, DOG1 (33), and positive regulation of germination, PELPK1 (34) (See Additional file 2: Table S2). Only two *sylvestris* unique genes had annotated functions (Additional file 2: Table S2). One corresponds to GSH-induced LITAF, which negatively regulates hypersensitive cell death in *Arabidopsis* (35). The other one corresponds to FAR1 (far-red-impaired response) related sequence, with roles in diverse developmental and physiological processes (36,37).

In addition, we assembled the plastid and mitochondrial genomes of the cultivar Farga, which were not provided as separate assemblies in the previous release (22) (see materials and methods). The final assembly of the plastid genome comprised 155,658 base pairs (bp) (Additional file 2: Table S3), in agreement with previously reported olive plastid sequences, which range from 155,531 to 155,896 bp (16,38,39). We annotated 130 genes out of the 130-133 genes reported for other olive plastid genomes (see Additional file 1: Fig. S2, Additional file 2: Table S3) (16,38), of which 85 are protein coding genes, 37 are transfer RNAs, and eight are ribosomal RNAs. The final assembly of the mitochondrial genome has a size of 755,572 bp (Additional file 2: Table S3), which is similar to that of previously sequenced wild olive mitochondrial genomes (710,737 – 769,995 bp) (39). The coding regions in the olive mitochondrion comprise 46 protein-coding genes, 3 ribosomal RNA genes, and 26 transfer RNA genes (Additional file 1: Fig. S3, Additional file 2: Table S3).

### Contrasting genetic diversity patterns in organellar and nuclear genomes

We used this improved reference genome assembly (Oe9) to call SNPs of all individuals at the nuclear, plastid, and mitochondrial genomes. Altogether, for wild and cultivated olives (subsp. *europaea*), we obtained a total of 16,604,110 polymorphic positions uniformly distributed along the nuclear genome (Additional file 1: Fig. S4), 81 in the plastid genome, and 2,465 in the mitochondrial genome (see Additional file 1: Fig. S2, S3). In the plastid genome, a large region (~25 Kb) was found to be fully conserved and devoid of SNPs in all analyzed individuals (Additional file 1: Fig. S2). Interestingly this region includes the largest plastid gene, ycf2, which has also been found to exhibit low rates of nucleotide substitution in other plants (40). This gene is essential for plant survival, however its exact function is unknown (41,42). This conserved region also comprises other genes, including ycf15, rps7, rps12, ndhB, rRNA and tRNA. All individuals presented similar amounts of nuclear polymorphisms relative to the Oe9 reference (Fig. **1a**). Interestingly, the *sylvestris* from northern Spain (*sylvestris*-S) has a higher number of homozygous SNPs and a lower number of heterozygous SNPs (Fig. **1a**), suggesting a historically small population size. In contrast, the recently published *sylvestris* from Turkey (*sylvestris*-T) has a number of homozygous and heterozygous SNPs in the range of variation found in cultivars (Fig. **1a**). This observation was also obtained when the SNP calling was performed using *sylvestris-T* genome as a reference (data not shown).

**Figure 1.**
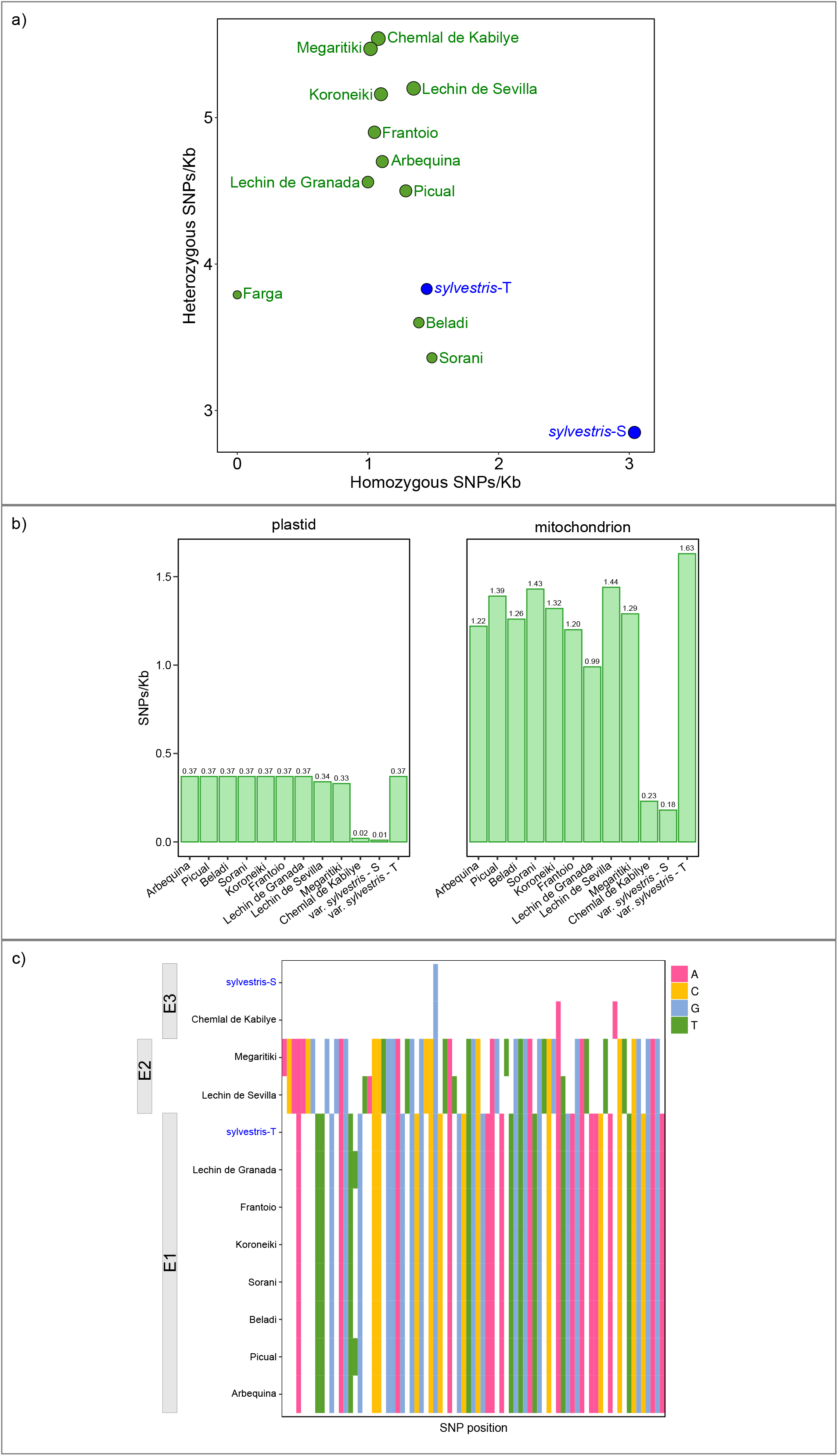
SNP density (SNPs/Kb) in sequenced individuals. **(a)** Homozygous versus heterozygous SNPs for each accession. Dot size correlates with the total amount of SNPs. All the cultivars are marked in green and var. *sylvestris* in blue. **(b)** SNP densities for the plastid and mitochondrial genomes. **(c)** Plot showing the relative position and identity of plastid SNPs as compared to the cultivar Farga (reference genome). Bars on the left show the main plastid haplotypes of the individuals as described by Besnard et al., 2011, 2007.

Strikingly, the patterns of polymorphisms in the organellar genomes did not follow the gradient described above for the nuclear genome (Fig. **1b**). In contrast to the nuclear genome, the plastid and mitochondrial genomes of the wild individual *sylvestris*-S and the cultivar Chemlal de Kabilye from Algeria show a notably lower number of SNPs relative to the Farga reference genome (Fig. **1b**). Plastid genomes can be arranged into three groups based on nucleotide polymorphisms, which are congruent with the three main plastid lineages (E1, E2, and E3) already described (Fig. **1c**) (16,43). More interestingly, the *sylvestris-S* shares the same plastid haplotype as cultivars Chemlal de Kabilye and Farga (E3), having only one SNP difference when compared with the latter cultivar (Fig. **1c**). Hence, the nuclear and organellar genomes presumably reflect different evolutionary histories. Notably, the organellar genomes of the cultivars Farga, Chemlal de Kabilye, and the wild individual *sylvestris*-S sequenced in our study show a very close genetic relationship (Fig. **1b,c**). As organelles are maternally inherited in olive (44), our results suggest that these cultivars and *sylvestris*-S share a most recent common maternal ancestor. Similarly, ‘Megaritiki’ (from Greece) and ‘Lechin de Sevilla’ (Spain) share the same plastid haplotype (E2), which is different to that of all the other cultivars. Interestingly, this haplotype can also be found in wild populations but exclusively from central and western Mediterranean areas (17).

Furthermore, we found a very close relationship at the organellar and nuclear levels between the eastern cultivars ‘Sorani’ and ‘Beladi’ -from Syria and Lebanon- and ‘Picual’ and ‘Lechín de Granada’, two of the most important cultivars of the south of Spain. This finding provides support for a recently proposed hypothesis of a recent bottleneck affecting a subset of western cultivars (12). This ‘local’ bottleneck affected only western olive cultivars (mostly from southern Spain and Portugal) and it was probably related to the introduction of olive germplasm into southern Spain during the Muslim period. This period began c. 700 AD, lasted eight centuries and possibly reshaped the cultivated olive germplasm of the Iberian Peninsula due to purported migration of cultivars from the Levant.

### Demographic analysis supports a population bottleneck coupled to the early domestication period

The average pairwise nucleotide diversity based on 20 Kb windows along cultivar genomes (4.48×10^-3^) is slightly lower than that of wild forms (5.54×10^-3^). This is consistent with previous studies based on inter-simple-sequence-repeats (ISSRs) (3), allozyme polymorphisms (45), simple-sequence repeats (SSRs) (12), and plastid DNA variation (16). Similarly, a recent transcriptome-based analysis reported slightly lower genetic diversity in cultivars compared to wild olives, leading to the suggestion of a weak-moderate population bottleneck during domestication (11).

In general, lower genetic diversity in cultivars is commonly associated with genetic bottlenecks during domestication (46). The difference between wild and cultivated trees observed here is less pronounced than that of many domesticated plant species (46,47). However, perennial crops often do not show evident domestication bottlenecks, in part because vegetative propagation means that perennials may not be many generations removed from their ancestral genetic diversity (48). In order to explore this possibility, we inferred the demographic history of olive using SMC++ v1.15.2, a tool that handles unphased genomes (49) (see material and methods). The results of this analysis show evidence for a continuous decline in population size starting approximately 11 kya, until ~4.5 kya (Additional file 1: Fig. S5). The end of this period corresponds to the olive domestication frametime (~6.0 kya) (8) and implies a possible domestication bottleneck. Interestingly, since olive domestication, an expansion of effective population size (Ne) can be observed. Altogether, these results suggest a mild bottleneck followed by a sustained population expansion during olive domestication.

### Population structure and phylogenetic relationships of the olive tree

We reconstructed the phylogenetic relationships of wild and cultivated olives using nuclear, plastid and mitochondrial SNPs separately. In addition, we tested introgression using a model-based genetic structure analysis (see material and methods). Interestingly, the nuclear polymorphisms based phylogeny (Fig. **2a**) places *sylvestris-T* intermingled within cultivars, and close to ‘Beladi’ and ‘Sorani’, which are main cultivars from Lebanon and Syria, respectively. This result remained the same when *sylvestris*-T was used as reference for the SNP calling (data not shown). Based on this and further results discussed in the following sections, we suggest that *sylvestris*-T represents a feral individual and may have been misidentified. Considering the results above, this might have contributed to apparent lower differences between wild and cultivated olives in terms of genetic diversity. Genetic structure analyses including all thirteen individuals from subsp. *europaea* suggested the presence of three distinct ancestral genetic clusters, which are differentially present among the different individuals (Fig. **2b**, Additional file 2: Table S4). Based on this, we distinguished three groups: one composed only by cluster 1 (‘Beladi’, ‘Sorani’), another composed by a mixture of cluster 1 and 2 (‘Lechin de Granada’, ‘Picual’), and the largest consisting of a mixture of the three clusters (‘Farga’, ‘Arbequina’, ‘Koroneiki’, ‘Frantoio’, ‘Lechin de Sevilla’, ‘Megaritiki’, ‘Chemlal de Kabilye’). When we analyzed the two oleaster samples, we observed that *sylvestris*-S is composed only by cluster 3, while *sylvestris*-T is composed largely by cluster 1, similar to ‘Beladi’ and ‘Sorani’, but with a small fraction of cluster 2. Interestingly all cultivars shared cluster 1, which is pervasive and enriched in cultivars from the eastern MB in different proportions, suggesting that this may be a consistent fingerprint of the primary domestication event that took place in this area (17). The dominance of this genetic background among cultivars suggests that the cultivars derive from a common primary domestication process. However, the presence of additional gene pools within the cultivars depicts patterns that could have resulted from preferential selection of genetic variants among standing variation, from separate domestication events, or from introgression events with wild populations.

**Figure 2.**
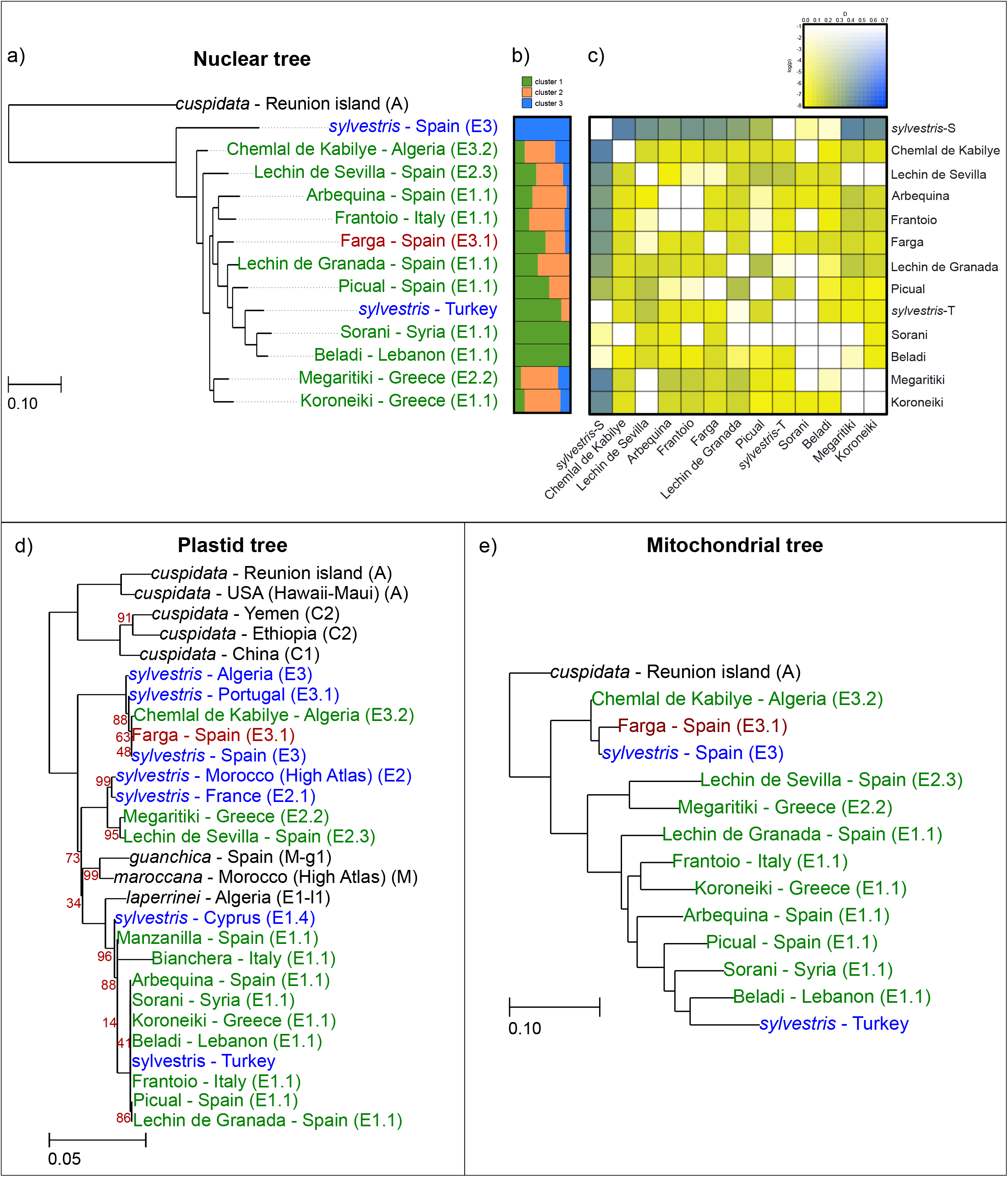
Maximum likelihood species tree derived from the SNPs data: **(a)** Nuclear phylogeny. Cultivated olives are shown in green and wild olives in blue. The geographical location of the accession and the plastid haplotype are indicated. Only bootstrap values below 100% are indicated. **(b)** Bayesian clustering for the nuclear SNP data estimated in Structure v2.3. Structure bar plot shows the genetic clusters differentiated by color. **(c)** Heatmap showing the *D*-statistic and its p-value. Blue color indicates higher *D*-statistics, and more saturated colors indicate greater significance. **(d)** plastid phylogeny. The colors and characteristics are the same as in (a). **(e)** mitochondrial phylogeny. The colors and characteristics are the same as in (a).

To reconstruct the plastid tree, we included the available plastid genomes of four additional accessions of *O. europaea* subsp. *cuspidata* and one individual of other subspecies that are publicly available (see material and methods). In this phylogeny (Fig. **2d**) *cuspidata* individuals grouped together in congruence with previous results (16,43). The topology of the phylogenetic tree does not change if we use other plastid reference genomes such as Manzanilla (NCBI:FN996972), ‘Bianchera’ (NCBI: NC_013707), or ‘Oeiras 1’ (NCBI: MG255763) for the SNP calling (data not shown). Remarkably, consistent with the polymorphism patterns discussed earlier, we observed incongruencies between the organellar and nuclear trees (Additional file 1: Fig. S6), which gives support for a phylogenetic signal of hybridization (50–52). In particular, all cultivars (plus the possible feral *sylvestris*-T) are monophyletic in the nuclear tree (being *sylvestris-S* the earliest branching), whereas the plastid tree shows that cultivars are polyphyletic, falling into three independent clades, each of them associated with different *sylvestris* samples. Moreover, attending to the position of samples from other subspecies (such as *guanchica, marrocana*, and *laperrinei*), which are intermingled within the subspecies *europaea*, the plastid genome suggest a polyphyly of the subspecies *europaea*.

A particularly strong phylogenetic incongruence observed among cultivars involves the cultivars Farga and Chemlal de Kabilye, which cluster together with the other cultivated olives in the nuclear tree, but are sister to var. *sylvestris* from Spain, far from the other cultivars in both plastid and mitochondrial trees (Fig. **2d,e**). This result supports previous evaluation of maternal inheritance of plastid and mitochondria (21). Particularly, these results suggest that the maternal line of cultivars Farga and Chemlal de Kabilye originated from western Mediterranean wild olives (which carry the E3 plastid haplotype), while the paternal line originated mostly from domesticated individuals from the eastern Mediterranean basin. A similar pattern has been observed in a previous study that combined a large sample of cultivated olives and oleasters, in which most cultivars were assigned to the eastern genetic pool, even those with maternal lineages that originated from the western Mediterranean basin (11,17). All these results reinforce the idea that cultivars are either from the eastern genetic pool or admixed forms (6,10,17), and support secondary admixture processes in the western Mediterranean basin, with contribution from western populations of var. *sylvestris*, as clearly shown by the plastid lineages. Indeed, the largest wild populations of olives are found in the western Mediterranean.

In sum, the plastid tree suggests genetic contributions, at least in the maternal lineage, of three different genetic pools that may be in agreement with multilocal domestication processes, whereas the nuclear tree suggests a dominant, congruent signal shared by all sequenced cultivars that is consistent with a unique primary domestication center in the eastern MB.

### Genetic introgression from western var. *sylvestris* genetic pools among cultivated olives

Previous phylogenetic analyses have suggested the existence of ancestral inter-subspecies hybridization in deep nodes of the evolutionary tree of *O. europaea* (25). Our population genomics results discussed above and a recently published transcriptome-based population analysis (11) suggest recent intraspecific genetic admixture between western cultivars and western *sylvestris*. To incorporate genetic admixture into a phylogenetic framework, we reconstructed a split network tree based on nuclear genome data (Fig. **3**), which revealed a heavily reticulated structure mostly affecting the relationships among all olive samples. In particular, most samples with the exception of cultivars Sorani, Beladi and *sylvestris-T* appear in a heavily reticulated area that shows a distant connection with *sylvestris*-S.

**Figure 3.**
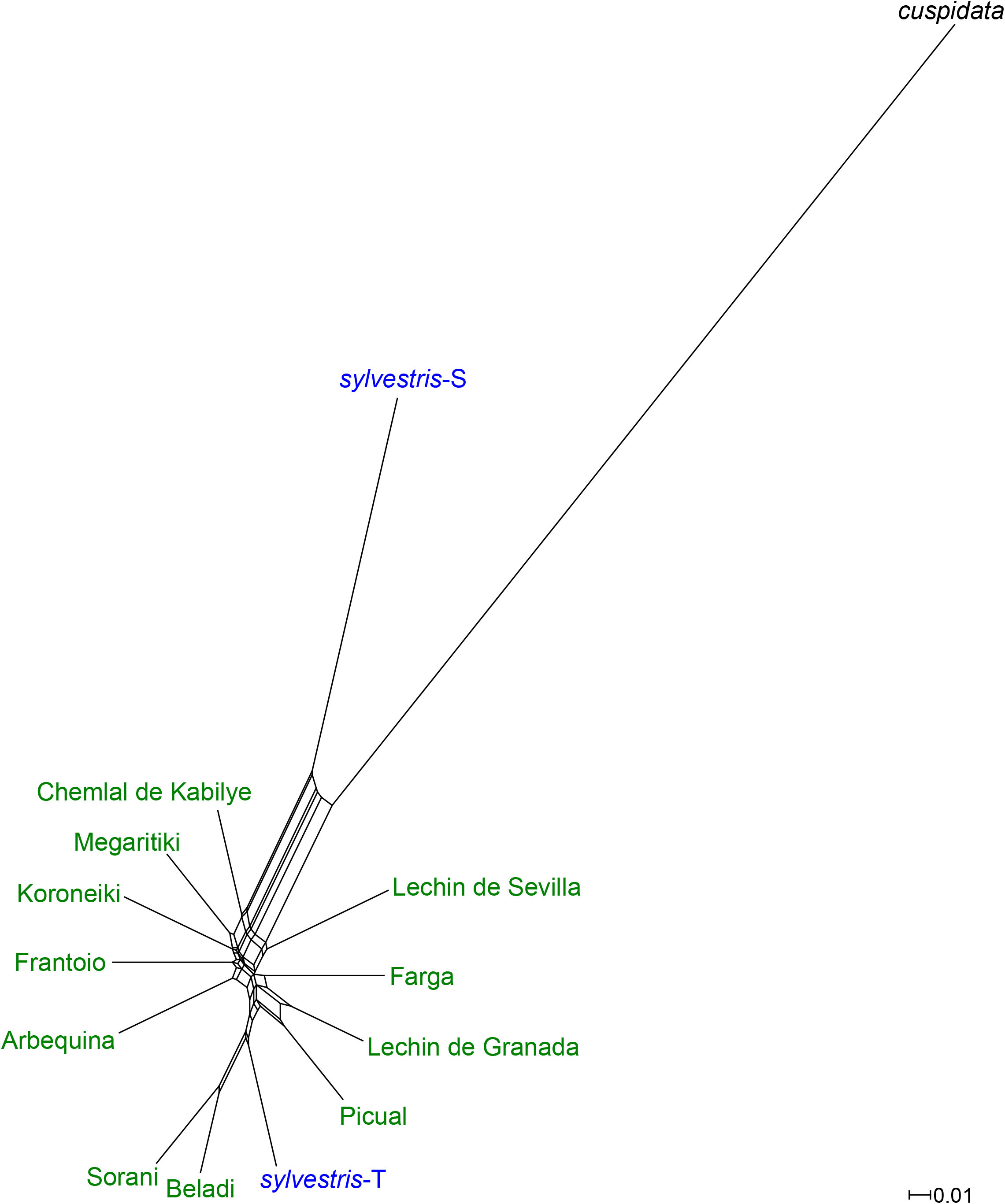
SplitsTree derived from nuclear SNPs. All the cultivars are marked in green and var. *sylvestris* in blue. The neighbor-net method is used here to explore data conflict and not to estimate phylogeny.

Consistent with the population genomics results, *sylvestris*-T appears well-embedded within cultivars and with little signs of reticulation. Overall, such a reticulated network can be the result of hybridization (including introgression) or incomplete lineage sorting. In order to detect evidence of introgression among the olive samples, we ran the ABBA-BABA test (53) using the program Dsuite v0.1 r3 (54) (see material and methods) for all quartets of the phylogenetic tree (Fig. **2a**). This analysis shows support for introgression between *sylvestris*-S and the lineages of ‘Chemlal de Kabilye’, ‘Megaritiki’, ‘Koroneiki’, with a significant D-statistic around 0.43 (Fig. **2c**). In addition, there seems to be some support (D-statistic ~0.35) for introgression between *sylvestris-S* and lineages of ‘Farga’, ‘Arbequina’, ‘Frantoio’, ‘Lechin de Sevilla’, and ‘Lechin de Granada’. When we investigated the D-statistic over sliding windows across the genome (see material and methods) using the eight individuals with introgression we found a different amount of genomic regions with a D-statistic higher than 0.5. Individuals with higher D-statistics have larger genomic regions with signatures of introgression (Additional file 1: Fig. S7a). Then, we assessed whether specific functions were enriched in genes located within introgressed regions and found no significant enrichment.

In agreement with the split-tree, ‘Sorani’, ‘Beladi’, and *sylvestris-T* show no signs of introgression with *sylvestris*-S. This contrasts with the other analyzed cultivars that have been strongly introgressed from wild olives of the western MB (here represented by *sylvestris*-S). Importantly, the level of introgression is largely independent of the plastid haplotype (all cultivars of plastid haplotype E2 and E3, and some of E1 show signs of introgression), suggesting that, in at least some cases, different introgressions have occurred in the maternal and paternal lineages, independently. Most of the introgressed cultivars originate from areas of the western or central MB. The three non-introgressed genotypes ‘Sorani’, ‘Beladi’, and *sylvestris*-T were all sampled from the eastern MB (Lebanon, Syria and Turkey), close to the purported origin of olive primary domestication. Overall, our genome-based results are consistent with earlier results on a broader dataset and on transcriptome-based analysis (11).

### Identification of genes under selection

Olive trees were domesticated for their fruits, either as a source of oil or edible fruits (55). Genes associated with agronomic traits may have undergone positive selection during domestication. In order to search for genes putatively under positive selection in the cultivars, we first classified the SNPs into intergenic, intronic, and coding. We further classified coding SNPs according to whether they imply synonymous or nonsynonymous changes (see material and methods). Additional file 1: Fig. S6 shows that there is a higher percentage of SNPs in intergenic regions (4.7 SNPs/Kb), followed by intronic (0.8 SNPs/Kb) and coding regions (0.3 SNPs/kb). Moreover, the number of synonymous and nonsynonymous changes is similar across samples. In order to assess selection we first measured the ratio of nonsynonymous and synonymous nucleotide diversity (*π*N/*π*S) in all the individuals included in this study. All had similar *π*N/*π*S values, with an average of 0.40 (Table 2), suggesting similar strengths of selective pressure across all sequenced individuals. This ratio is similar to that found in other trees such as *Populus nigra* (0.48) (56) and *Populus trichocarpa* (0.40) (57).

**Table 2.**
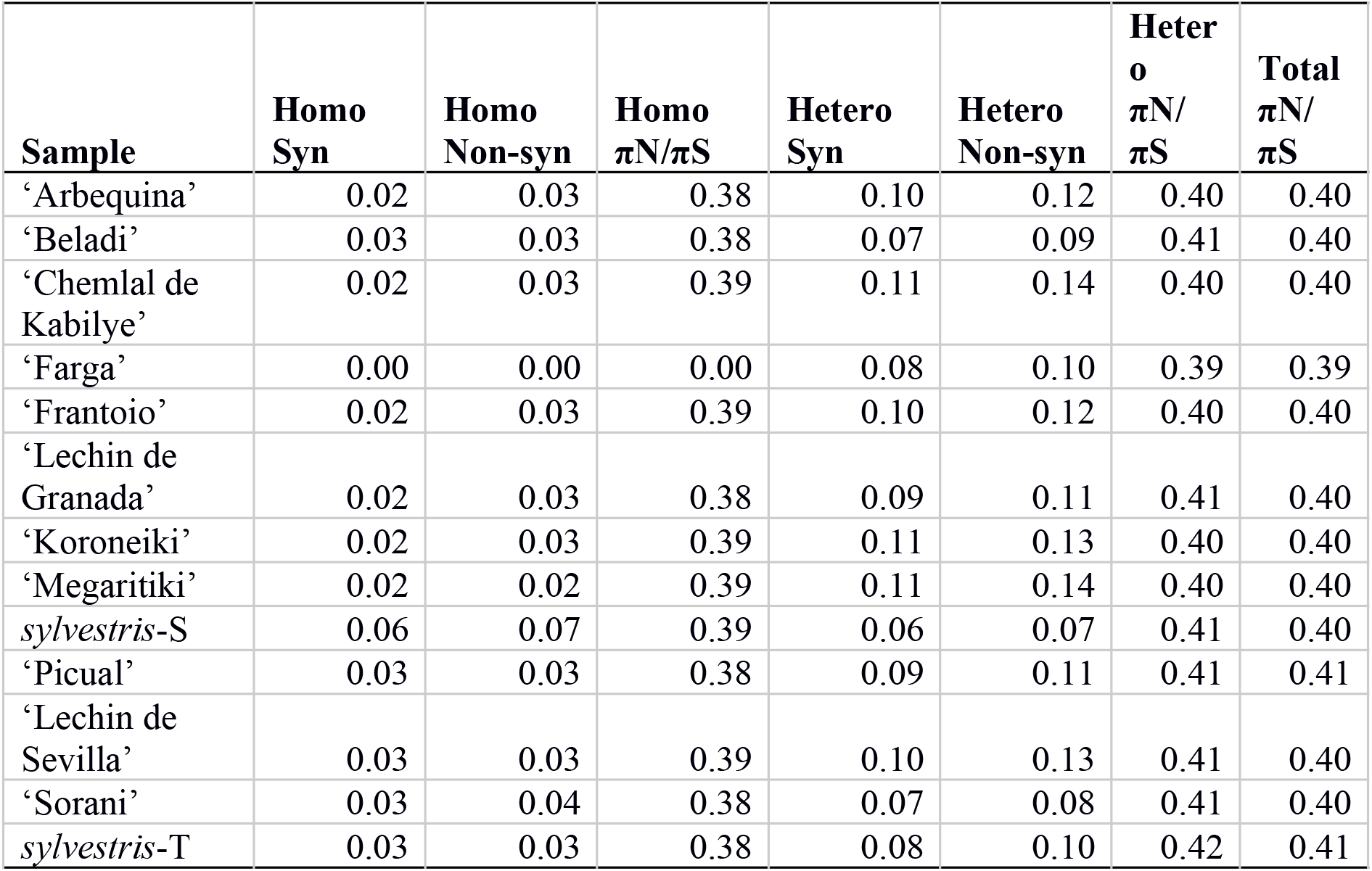
Number of synonymous and nonsynonymous homozygous SNPs, and synonymous and nonsynonymous heterozygous SNPs per individual. Columns in order show: number of homozygous-synonymous SNPs, number of homozygous-non-synonymous SNPs, *π*N/*π*S ratio of homozygous SNPs, number of heterozygous-synonymous SNPs, number of heterozygous-non-synonymous SNPs, *π*N/*π*S ratio of heterozygous SNPs, *π*N/*π*S ratio of total number of SNPs.

When we analyzed the SNPs that can produce a nonsynonymous change, including heterozygous and homozygous SNPs, we found that a total of 32,723 proteins (59% of the predicted Oe9 proteome) have at least one SNP with nonsynonymous change, of which 14,985 are common for all the individuals (Additional file 2: Table S5). On the other hand, the list of proteins that did not show any nonsynonymous change were used to perform a functional enrichment analysis, and we found overrepresented GO terms associated with molecular functions, cellular components, and one biological process (see Additional file 2: Table S6). These functional categories may be under a particularly strong purifying selection. When we look at the genes without SNPs, 21,492 genes are present in all individuals and few are unique to each individual (Additional file 2: Table S5). Interestingly, *sylvestris-S* has the largest number of genes (1,581) without SNPs relative to cultivar Farga.

To further determine whether one or more proteins are under positive selection in cultivated olives, we searched for evidence of recent selective sweeps by quantifying site frequency spectrum (SFS) deviations relative to genome-wide patterns using the composite likelihood ratio (CLR) statistic implemented in SweeD (58) (see material and methods). A total of 555 regions, which fell within the top 0.5% of the observed distribution were considered as candidates for positive selection. After merging the overlapping regions (see material and methods), we kept only 181 genomic regions, from which 5% were larger than 50 kb and 56% shorter than 10 Kb. A total of 84 genes were identified in 69 sweep regions. Among the genes found within sweep regions, some are associated with lipid, carbohydrate, and amino acid metabolism, and others with stress tolerance, solute transport, and RNA processing (Additional file 2: Table S7). Remarkably, we did not find any gene related to fatty acid metabolism and accumulation, although this has presumably been one of the most important characters under selection in olive domestication. Moreover, a recent study found 19 genes associated with 5 important agronomic traits in olive (59) and despite 14 of them being found in Farga (see Additional file 2: Table S8), none was present in the detected selective sweep regions. Similarly, a study based on transcriptomes of 68 different wild and cultivated samples using statistical approaches (PCAdapt (60) and BayeScan (61)) failed to identify candidate genes under selection associated with oil content or fruit size (11). They did, however, detect ten genes as strong candidates for selection that were associated with transcriptional and translational activities and to the cell cycle. Also, it was proposed that domestication in olive may be more related to changes in gene expression than changes in protein function in agreement with its evolutionary history (11). Further studies with more cultivated and wild samples will be needed to test this inference further.

Since some cultivars show signature of introgression with *sylvestris-S*, we assessed whether adaptive introgression has contributed to olive domestication. When we analysed the 181 selective sweep regions, we found that on average 31% of these regions were overlapping introgressed regions (Additional file 1: Fig. S7b, Additional file 2: Table S9), with ‘Megaritiki’ being the cultivar with more common regions (51%), followed by ‘Chemlal de Kabilye’ (45%). Overlapping regions of introgression with selective sweeps suggest that adaptive introgression seems to have played an important role in olive domestication, as it has been observed in other crops (62).

## Discussion

### Improved assemblies for nuclear and organellar genomes

Our first reference genome assembly for *Olea europaea* (Oe6 version) (22) provided a needed resource for researchers interested in understanding the genetic basis of olive traits and the process of domestication. Here we improved the assembly (Oe9) by anchoring it to chromosomes using a publicly available genetic map (24). In addition, we produced individual annotated assemblies for the plastid and mitochondrial genomes, which were not provided as separate assemblies in earlier releases. Our comparison of our improved nuclear assembly with the recently released assembly of an oleaster from Turkey (23) and the identification of shared duplicated regions provides additional support to the previously proposed ancient polyploidization in olives (25). Unexpectedly, we found apparent differences in gene content even using a conservative approach that is independent of differences in annotation methodologies. These differences could reflect shortcomings of the assemblies, changes occurred during domestication, or variation in gene content among individuals, as observed in grapevine, another perennial crop (63). Further efforts in improving reference assemblies for the olive tree with new long read technologies will definitely enhance our understanding on the effect of genome rearrangements, including deletions of genomic regions during the evolution of olive.

### Reliable genetic sources: feral individuals vs. true wild trees

Understanding the domestication process in olives requires careful comparison between reliably identified cultivars and wild individuals. Indeed, plant choice is crucial before starting any genomic studies. In order to use reliable genetic sources, biology features of olive germplasm have to be considered in the field: a) high phenotypic plasticity and resilience; b) long reach of aerial transported pollen; c) the ability to cross between genetically distinct individuals; and d) the long lifespan of the trees (often reaching several centuries and millennia). These characteristics make it extremely difficult to tell apart pure wild individuals from those highly introgressed or even cultivars (64). As stressed earlier, we took great care in selecting an individual from an isolated wild olive population, which had been studied over the last years and meets key characteristics that conform to a true oleaster. Our phylogenetic analysis based on the nuclear genome places this wild individual at the earliest branching position within the individuals of the subspecies *europaea*, sister to the rest, which is fully consistent with its ascription to a wild western Mediterranean genetic pool. In stark contrast, the results for the wild sample from Turkey (23) were puzzling. The phylogenetic analysis placed it at a relatively shallow clade, forming a tight cluster with two cultivars from the eastern MB. This result could be explained through four hypotheses: i) the Turkish individual identified as *sylvestris* is actually a feral olive, ii) this tree is an oleaster highly introgressed with cultivars; iii) considering that the cultivated olive originated from wild oleasters, this pattern may represent one of multiple primary domestication events; or, alternatively (iv) a separate, very recent domestication event. However, hypotheses (iii) and (iv) are at odds with the observed topology in which the well-defined clade formed by *sylvestris-T*, ‘Beladi’, and ‘Sorani’ is placed at a rather shallow position of the tree, embedded within a larger clade of other cultivars; whereas sister-clade relationships to other cultivars would be expected from independent domestication events. Further population analysis, including the search for introgression signals discard hypothesis (ii), as *sylvestris-T* individual showed weak signature of introgression and instead a genetic structure highly similar to the nearby sampled cultivars. In addition, the rarity of wild olive trees in the eastern Mediterranean (65) together with the small number of phenotypic characters to differentiate cultivated olives from oleaster, must be considered. The “Flora of Turkey” (page 156) states that “… spontaneous seedlings may revert to var. *sylvestris”* which, despite the misuse of the term reversion, highlights the phenotypic similarity between ferals and true oleasters in the region (65). Earlier studies at a large geographical scale already mentioned the difficulty to find genuine oleasters in the eastern MB (45,66). Recent experiences confirmed this fact, analyzing putative wild olives from Turkey (15) and Israel (12) that appeared to be feral after genetic analyses. Given all these facts, we suggest that *sylvestris-T* is a feral individual, and we have treated it as such to avoid confounding conclusions. Given the open debate on possible alternative centres of domestication, and the interest in tracing the genetic aftermath of domestication in olives, the meticulous choice of additional wild populations will be necessary. In order to reliably reconstruct complex domestication processes, we stress the importance of careful sample collection of olive trees in the field using ecological (67) and key morphological characters (68), assisted with molecular screening of olive material, before undertaking genome sequencing of wild trees.

### Genomics support for primary and secondary events of domestication driven by introgression from wild genetic pools

This study shows the first phylogenetic analysis of genome-wide sequences of the Mediterranean olives. Our results, together with those from previous analyses (11), suggests that cultivated individuals have slightly lower nucleotide diversity as compared with wild individuals. However, our demographic analyses support the existence of a relatively small population size at the time of domestication, with a steady decrease in population size preceding domestication as has been inferred for some other crops (69,70). These analyses are consistent with earlier studies suggesting a narrow distribution (and hence limited population size) of oleaster populations in the eastern MB over the last 150,000 years (8). The demographic analysis also indicates a mild population bottleneck around 5000-7000 years ago, consistent with the proposed period for primary domestication of olives in the Levant around 6000 years ago (8). Interestingly, our analysis also suggests a rapid increase in population size following the domestication bottleneck, likely coupled to the expansion of olive cultures in broader Eastern Mediterranean countries. Altogether, these results suggest that one ancestral genetic pool, likely deriving from a founding population from eastern wild trees, is pervasive among cultivars. This is consistent with common ancestry at a primary domestication event from which all cultivars descend.

However, a common primary domestication event is not incompatible with subsequent nested domestication processes, perhaps driving the adaptation to local conditions or the selection of specific traits. Considering our results and previous ones from numerous molecular techniques, the emerging pattern suggests a scenario of multiple events of hybridization between individuals descending from this primary domestication event ameliorated with genes from different wild populations. These processes have affected cultivars from the central and western MB to different degrees. We propose that extensive gene flow between genetically rich wild and cultivated olives occurred through the expansion and diversification of olive crops by ancient Mediterranean cultures (Phoenicians, Romans, Arabs) (71). It is likely that these introgression events, followed by artificial selection of desired characters, resulted in the incorporation of alleles from wild populations and facilitated the creation of specific olive cultivars adapted to local environments.

The origin of the introgressed genetic material can be better inferred when the donor lineage can be traced back through the maternal line. Earlier studies have shown the presence of clearly distinct haplotype groups (E1, E2, and E3) among cultivars (16,43). E1 is present in wild trees from the eastern MB and it is likely the signature of the primary domestication event, since it is shared by the 90% of the current olive cultivars (17). This is particularly true for individuals sampled from regions close to the origin of domestication, ‘Sorani’ (Syria), and ‘Beladi’ (Lebanon) and the feral individual from Turkey, which show little or no signs of introgression. Cultivars carrying the haplotypes E2 and E3, also found in wild forms of the western MB, were those often revealing a blueprint of introgression in our analysis.

Particularly, the cultivars Farga and Chemlal de Kabilye have a different phylogenetic history than that of the other cultivars suggesting secondary hybridization of cultivars with wild oleasters from the Iberian Peninsula, in which a lineage similar to var. *sylvestris* from Spain acted as the maternal line. Consistent with this, the nuclear genome of these two cultivars shows signs of introgression with var. *sylvestris* from Spain. However, this introgression signal is also detected in other cultivars, irrespective of the plastid haplotype lineage. Altogether, these results suggest that admixture with wild individuals from western populations of var. *sylvestris* has been common, and has taken place multiple times, both in the maternal and paternal lineages.

Based on plastid haplotypes of cultivars, introgression from at least two different wild genetic pools, other than the one involved in the primary domestication, has contributed to the nuclear genomes of cultivars, affecting different subsets of cultivars. These different genetic pools are highly divergent from each other, as shown in our phylogenetic reconstructions from plastid genomes, which suggest that the subspecies *europaea* might have multiple origins (polyphyletic). Some other studies with plastid and nuclear markers have shown similar results (2,3,43). Despite the detectable signal of introgression in the nuclear genomes, all cultivars, including introgressed ones, are monophyletic in our reconstructions (considering *sylvestris-T* as a feral). This is also the case of the cultivar Farga, which shares a maternal ancestor with the *sylvestris-S* individual analyzed here. These two individuals are closely related when organellar genomes are used for phylogenetic reconstruction, but cultivar Farga appears well embedded within the cultivars clade in the nuclear genome phylogeny. This indicates that the introgressed material in the nuclear genome has been reduced through subsequent crosses following the initial hybridization. Finally, the closest similarity of the two cultivars of southern Spain ‘Picual’ and ‘Lechín de Granada’ with the eastern cultivars ‘Sorani’ and ‘Beladi’ support a previously proposed secondary migration of eastern genetic pools possibly during the Muslim period of the Iberian Peninsula or even earlier (12).

Altogether, the phylogenetic and admixture analysis show that rampant hybridization has shaped the evolutionary history of the different lineages of olives. We thus hypothesize a recurrent pattern of domestication that involves a primary domestication event in the east of the MB, followed by nested secondary (and tertiary, and so on) local domestication events with the involvement of additional wild genetic pools, forming a domestication continuum (6). Admittedly, increased genomic sampling, particularly of true wild populations, is needed to help describe these complex patterns of evolution in *Olea europaea* across the numerous areas of the Mediterranean basin.

### Lack of a clear domestication syndrome

Cultivated olives have undergone a complex domestication process, which has led to morphological and physiological changes. The main traits selected during the transition between oleasters and cultivars are fruit size and oil content (13,55,68). The domestication scenario described in the previous section, which is punctuated by hybridization, may make it difficult to detect genes selected during the process of domestication. We used branch-site models to detect sequence changes likely associated with selection. We detected very few genes putatively selected at the lineage subtending all sequenced cultivars. However, when analyzing each cultivar lineage individually some patterns emerged. In particular, genes positively selected in cultivated olives were associated with a response to biotic and abiotic stress. However, further analyses are needed to ascertain whether they indeed play a role in trait selection during domestication (see (72) for secondary compounds). Previous analyses detected few signs of positive selection affecting protein-coding regions, but they did detect differences in expression levels between cultivars and wild individuals for specific genes (11). This led to the conclusion that selection may have acted on non-coding regions that drive gene expression. However, given the difficulty of controlling for other factors driving gene expression (different periods of the year, local environmental conditions), we believe this result must be viewed with caution. Alternative approaches are needed to detect alleles whose presence in the cultivars were selected through domestication. We cannot rely on models that assume only vertical evolution, but would rather search for shared conserved or introgressed regions across different cultivars sharing similar phenotypes (68). To be effective, a much larger sampling of genomic sequence will be needed for such an approach. In addition, assemblies from representative lineages of reliable wild olives will help to better trace the origin of different introgressed regions in the genomes of cultivars. Overall, our new phylogenomic and genetic analyses of whole-genome sequences show evidence for a complex process of reticulation disrupting historical isolation in the course of olive domestication.

## Material and Methods

### Scaffolding of the cultivated olive genome using a linkage map

A new, improved version of the *O. europaea* genome assembly (Oe9) was produced by anchoring the Oe6 version (22) to a publicly available genetic map (24) using ALLMAPS (73). First, we took the intersection of 7,042 markers for which a sequence was provided and mapped them with BWA v0.7.15 (74) to the Oe6 olive genome assembly. Filtering for minimum mapping quality 20 and fewer than 10 mismatches, we obtained 5,780 mappings. Intersecting these mapped markers with the 3,404 markers placed in the genetic map resulted in a set of 2,759 markers with both a genetic map position and an unambiguous physical location in the Oe6 assembly. ALLMAPS was then run with default parameters. A total of 2,362 markers were considered unique, of which 2,134 were anchored, and 228 unplaced.

### Functional annotation of the cultivated olive genome

In order to give more functional insight to our analysis, we decided to improve the functional annotation of the genes present in the Oe6 version of the genome by running Blast2go (75), which in turn ran a BLASTP (76) search against the non-redundant database (April 2018) and Interproscan (77) to detect protein domains on the annotated proteins.

### Comparison of the *europaea* and *sylvestris* genomes

To compare the two available genome assemblies of *O. europaea* subsp. *europaea* (var. *europaea* and *sylvestris*), we plotted their cumulative assembly lengths using the R Package ‘ggplot2’ (78). The predicted protein-coding gene sequences of the two assemblies were compared using a search with BLASTN (26). Results were filtered using cutoffs of 80% identity and e-value<1e-5. For the genes that did not have a hit, we analyzed whether they were covered by reads of the other sequenced accession. For this step we first mapped the reads of each genome against the other using BWA 0.7.6a-r433 (74). We considered a gene as individual-specific (e.g. *europaea*-specific) if it did not pass a filter of coverage >20 over 50% of the gene length in the other genome (e.g. *sylvestris*). With the aim of achieving a more conservative set of individual-specific genes, we applied a more stringent filter of coverage (>5). Finally, in order to search for the functions of the genes without a hit in the other genome (individual-specific), we performed a BLAST search against the NCBI non▯redundant database, and the same cutoffs as described before.

### Genome sequences

We sampled and sequenced twelve genomes of *O. europaea:* ten cultivars (‘Arbequina’, ‘Beladi’, ‘Picual’, ‘Sorani’, ‘Chemlal de Kabilye’, ‘Megaritiki’, ‘Lechin de Sevilla’, ‘Lechin de Granada’, ‘Frantoio’, and ‘Koroneiki’), one oleaster (*sylvestris*-S), and one subsp. *cuspidata* to be used as outgroup. These samples broadly covered the geographical distribution of the species in the MB (see Table 1). The authenticity of these cultivars was previously substantiated through molecular and morphological markers (9). The sequenced oleaster (referred to as *sylvestris*-S from now onwards) was collected in the North of Spain. All the samples were collected according to the local, national or international guidelines and legislation.

The DNA of all individuals was extracted as described in (22) and their genomes were sequenced using Illumina HiSeq 2000 paired-end technology to a sequencing depth ranging from 24 to 34x at the CNAG-CRG sequencing facilities, as described for the reference genome (22). In addition to these individuals, we used publicly available data of the reference genome sequence of the olive cultivar Farga, (22). ‘Farga’ is a cultivar from Catalonia (eastern Spain) with the E3.1 plastid haplotype and previously classified as representative of the central MB cultivated genepool (Table 1). We also used a recently published assembly of *O. europaea* var. *sylvestris* (referred as *sylvestris*-T from now onwards) (23), and downloaded fourteen plastid genomes of the Mediterranean olive from the NCBI database (see Table 1).

### Organelle Assemblies

The available reference genome sequence does not include separate scaffolds for mitochondrial and plastid genomes (22). Here, we assembled and annotated both organellar genomes of the cultivar Farga using paired-end (PE) and mate-pair (MP) data from the reference genome sequence project (22). Briefly, all genome shotgun Illumina reads were mapped using BWA v0.7.13-r1126 (74) to a reference plastid sequence (NC_013707), and a mitochondrial sequence (MG372119). Then reads were filtered allowing only those that mapped in proper pairs with a hard and soft clipping for a maximum of the 25% of the total length of the read. The plastid genome was assembled using NOVOPlasty v2.6.3 (79). The mitochondrial genome was first assembled with SPAdes v3.10.0 (80). Then, this initial assembly was scaffolded using SSPACE_Basic v2.0 (81) using PE and MP libraries. Finally, gaps were filled with GapFiller v1-10 (82).

The plastid and mitochondrial genomes were annotated by BLAST searches against previously annotated plastid (FN996972, MG255763) and mitochondrial (MG372119, KX545367) genomes. Gene structures (i.e. intron–exon boundaries) were defined using Exonerate v2.47.3 (83) using the “protein2genome” model. Annotations of tRNA genes were verified using tRNAscan-SE (84).

### Detection of Single Nucleotide Variants

To assess nucleotide diversity across sequences of the Mediterranean olive at the nuclear, plastid and mitochondrial levels, we called single nucleotide polymorphisms (SNPs) using the cultivar Farga genome as a reference for all the cases (for the nuclear genome we used the Oe9 version). Sequenced reads from each individual were mapped against the reference genome using BWA 0.7.6a-r433 (74), and SNPs were identified with GATK HaplotypeCaller v4.0.8.1 (85), setting ploidy to 2, and using thresholds for mapping quality (MQ > 40), quality by depth (QD > 2), Fisher strand bias (FS < 60), mapping quality rank sum test (MQRankSum > −12.5), read pos rank sum test (ReadPosRankSum > −8), strand odds ratio (SOR < 3), read depth of coverage (DP > = 10), and allelic depth (AD >= 3). Sites with missing alleles and spanning a deletion were also removed. Finally, VCFtools v0.1.15 (86) was used to filter out positions according to the number of alleles (--max-alleles 2), and minor allele frequency (--maf 0.04).

### Nucleotide diversity

Pairwise nucleotide diversity (*π*) was calculated using VCFtools v0.1.15 (86) per window of 20 Kb. The sequences were grouped together into two datasets: cultivated olives (‘Arbequina’, ‘Beladi’, ‘Picual’, ‘Sorani’, ‘Chemlal de Kabilye’, ‘Megaritiki’, ‘Lechin de Sevilla’, ‘Lechin de Granada’, ‘Farga’, ‘Frantoio’, and ‘Koroneiki’), and wild-feral olives (*sylvestris*-S and *sylvestris*-T).

### Demographic history of cultivated olives

To estimate population size histories in olive we employed SMC++ v1.15.2 (49), which is capable of analyzing unphased genomes. First we masked all regions larger than 5 Kb with a coverage < 10x in at least one of the samples included (‘Arbequina’, ‘Beladi’, ‘Picual’, ‘Sorani’, ‘Chemlal de Kabilye’, ‘Megaritiki’, ‘Lechin de Sevilla’, ‘Lechin de Granada’, ‘Farga’, ‘Frantoio’, and ‘Koroneiki’) using bedtools v2.26.0 (87). Then we ran SMC++ using default parameters and setting the T1, the most recent time point for population size history inference, to 10. Finally, a generation time of 20 years (12) and a mutation rate of 7.77 × 10^-9^ mutations per nucleotide per generation (88) were used to convert the scaled times and population sizes into real times and sizes.

### Admixture Mapping

Because of the large number of polymorphic positions in the nuclear genomes of the *O. europaea* samples, and computational limitations, we generated five subsets of 100,000 randomly-chosen polymorphic positions without overlaps, and analyzed them in parallel. Then we identified population structure without a priori grouping assumptions, using the Structure software v2.3.4 (89). Structure was run with 100,000 generations of ‘burn in’ and 100,000 Markov chain Monte Carlo (MCMC) iterations after burn-in for increasing K values ranging from 1 to 6, considering independent alleles and admixture of individuals. Simulations were repeated five times for each value of K. The optimal number of genetic clusters was determined using the ΔK method (90) using the software Structure Harvester (91). Finally, the optimal K value was visualized with DISTRUCT v1.1 (92).

### Phylogenetic analysis

Phylogenetic trees were reconstructed using SNP data from nuclear, plastid and mitochondrial genomes separately. In each case, the genome sequence of each sequenced individual was obtained by replacing the SNP positions in the respective reference genome, resulting in a pseudo-alignment of all the sequenced genomes. Specifically, for the nuclear dataset we included only homozygous SNPs. For the plastid genomes, we included additional sequences by aligning our genomes with the database genomes (see Table 1) using MAFFT v7.305b (93). All these alignments were trimmed using trimAl v1.4 (94) with the options -st 1 and -complementary, in order to remove all the non-informative positions. The final alignment had 11,774,982 variable positions for the nuclear genome, 325 for the plastid genome, and 2,731 for the mitochondrial genome. Phylogenetic trees were reconstructed from these alignments using RAxML v8.1.17 (95) and the GTR model as it is the most frequent evolutionary model found in previous studies. Support values were calculated based on 100 bootstrap searches using the rapid bootstrapping as implemented in RAxML. Additionally, for the nuclear data, we reconstructed a phylogenetic network using SplitsTree4 v4.14.5 and the NeighborNet approach (96).

### Analysis of Introgression with SNP Data

The ABBA-BABA test (53) was used to search for evidence of introgression among the olive samples. Dsuite v0.1 r3 (54) was employed to calculate the D-statistic from nuclear SNP data for all subsets of quartets that were compatible with the previously reconstructed phylogenetic tree (see above). For multiple hypothesis testing, we applied a false discovery rate correction to the p▯values (97). Then a heatmap showing the D-statistic and its p-value was plotted for all pairs of individuals using the plot_d.rb script (https://github.com/mmatschiner/tutorials/blob/master/analysis_of_introgression_with_snp_data/src/plot_d.rb). The Dsuite Dinvestigate tool was used in sliding windows of 5,000 SNPs, incremented by 1,000 SNPs to test whether the D-statistic is homogeneous or variable throughout the genome. It was tested in the trios with the strongest signals of introgression: Beladi - Chemlal de Kabilye - *sylvestris*-S, Beladi - Koroneiki - *sylvestris*-S, Beladi - Megaritiki - *sylvestris*-S, Beladi - Lechin de Granada - *sylvestris*-S, Beladi - Farga - *sylvestris*-S, Beladi - Lechin de Sevilla - *sylvestris*-S, Beladi - Arbequina - *sylvestris*-S, and Beladi - Frantoio - *sylvestris*-S.

### SNPs characterization

Nuclear SNPs were classified according to their genomic location as intergenic, intronic and coding. Coding SNPs were further classified into synonymous and nonsynonymous, according to the implied change in the respective codon. For the heterozygous positions, if at least one of the alleles was nonsynonymous we classified the position as nonsynonymous. GO term enrichment analyses of the proteins without nonsynonymous SNPs was calculated using FatiGO (98). To investigate the variation of nonsynonymous and synonymous SNPs within coding regions, we compared nonsynonymous changes per nonsynonymous site (*π*N) to synonymous changes per synonymous site (*π*S) using a synonymous: nonsynonymous site ratio of 1:3 (99).

### Identification of genes under selection

To detect protein-coding genes that have potentially undergone selection among the cultivated individuals we used an approach based on the site frequency spectrum (SFS). SweeD v3.1 (58) was used to identify selective sweeps in olive. This program is based on Sweepfinder (100) and uses a composite likelihood ratio (CLR) to identify loci showing a strong deviation in the site frequency spectrum (SFS) toward rare variants. We used as outgroup the subspecies *cuspidata* to infer ancestral alleles. SweeD was run separately for each scaffold and grid as the only parameter. The grid parameter was calculated per scaffold in order to have a measure of the CLR every 5 Kb (size of the scaffold/5000). Outliers were defined as the 0.5% with the most extreme p-values. Closer regions, with less than 10 bp distance were collapsed. Finally, a GO term enrichment analysis of the proteins that mapped to those regions was calculated using FatiGO (98).

## Ethics approval and consent to participate

Not applicable.

## Consent for publication

Not applicable.

## Availability of data and materials

The new version of the *O. europaea* var. *europaea* genome and the resequencing data produced in this project have been deposited in the ENA (European Nucleotide Archive) under the accession number PRJEB35540. The genome assembly and annotation are also available in https://denovo.cnag.cat/olive, together with a JBrowse genome browser and a Blast Server.

## Competing interests

The authors declare that they have no competing interests.

## Funding

This research has received funding from the European Research Council (ERC) under the European Union’s Horizon 2020 research and innovation programme (grant agreement No 724173; RETVOLUTION)”. IJ was supported in part by a grant from the Peruvian Ministry of Education: ‘Beca Presidente de la República’ (2013-III).

## Authors’ contributions

IJ, MMH, FC, JGG, TSA performed bioinformatics analysis. IJ and TG drafted a first version of the manuscript to which all authors contributed. IJ, MMH, FC, JGG, BSG, CMD, IGG, TSA, PV, TG read and approved the final version of the manuscript. TSA and IGG supervised the sequencing, assembly and annotation. CMD, BSG, PV provided samples, designed the selection of individuals and provided data and important scientific insights in the interpretation of the results. TG supervised the study.

## Acknowledgements

We want to acknowledge Guillaume Besnard, Manuel Nogales, Manuel Sánchez and Carlos Garcia-Verdugo for providing some of the plant samples used in this study. The authors want to thank Danelle Seymour and Rebecca Gaout for preparing some of the libraries. TG and PV acknowledge support from Banco Santander for the Olive Genome Sequencing project.

## Supplementary information

### Additional file 1

**Fig. S1. Genome comparison of *sylvestris* and *europaea*. (a)** Cumulative genome length per scaffold ranked in order of size for the genome assembly of *europaea* (red) and *sylvestris* (blue). A straight vertical line represents a perfect genome assembly. The horizontal plateaus indicates many small scaffolds. The top right end of each curve shows the total number of scaffolds. **(b)** Syntenic plot of the genome of *europaea* against *sylvestris* generated by SynMap.

**Fig. S2. Plastid genome of the cultivar Farga.** Protein coding genes are shown in green, rRNAs in light blue, and tRNAs in purple. The SNPs are shown per each individual included in this study in the following order starting from outside: ‘Arbequina’, ‘Picual’, ‘Beladi’, ‘Sorani’, ‘Koroneiki’, ‘Frantoio’, ‘Lechin de Granada’, ‘Lechin de Sevilla’, ‘ Megaritiki’, ‘Chemlal de Kabilye’, *sylvestris-S, sylvestris-T, maroccana, cerasiformis, guanchica, laperrinei, cuspidata-R, cuspidata-S, cuspidata-I*.

**Fig. S3. Mitochondrial genome of the cultivar Farga.** Protein coding genes are shown in green, rRNAs in light blue, and tRNAs in purple. The SNPs are shown per each individual included in this study in the following order starting from outside: ‘Arbequina’, ‘Picual’, ‘Beladi’, ‘Sorani’, ‘Koroneiki’, ‘Frantoio’, ‘Lechin de Granada’, ‘Lechin de Sevilla’, ‘ Megaritiki’, ‘Chemlal de Kabilye’, *sylvestris-S, sylvestris-T, maroccana, cerasiformis, guanchica, laperrinei, cuspidata-R, cuspidata-S, cuspidata-I*.

**Fig. S4. SNP distribution along the nuclear genome in windows of 100 Kb. (a)** homozygous SNPs, **(b)** heterozygous SNPs. Since cv. Farga was used as a reference genome, we do not expect homozygous SNPs for this sample.

**Fig. S5. SMC++ results for inferring population size histories in clutivated olives.** A generation time of 20 years was used to convert coalescent scaling to calendar time.

**Fig. S6. Number of homozygous and heterozygous SNPs (SNPs/Kb) in the intergenic, intronic and coding region of the genome.** The coding region was divided according to the changes that the allele can produce (synonymous and nonsynonymous).

**Fig. S7. a)** Plot showing the percentage of genomic regions with introgression (D-statistic >0.5). **b)** Plot showing the percentage of selective sweeps that are present in introgressed regions.

### Additional file 2

**Table S1.** General characteristics of the two genome assemblies of the cultivar Farga, and the genome assembly of the var. *sylvestris* from Turkey.

**Table S2.** List of unique genes of *europaea* and *sylvestris* with their homologous function.

**Table S3.** General characteristics of the plastid and mitochondrial genomes of the cultivar Farga.

**Table S4.** Admixture coefficient (Q) of each individual per cluster. This table was used to create the Fig. **2b**.

**Table S5.** Number of proteins with at least one nonsynonymous change per individual.

**Table S6.** List of the GO terms enriched in the list of proteins that do not have a nonsynonymous SNP. First column shows the term category, the second, the GO term, the third, the term level, the fourth, the p-value, and the fifth, the term name.

**Table S7.** List of proteins in regions with selected sweeps and their associated function.

**Table S8.** Blast results of the 19 genes of sylvestris-T against Farga. The results were filtered by %identity >90 and e-value<1e-5.

**Table S9.** Selective sweeps present in introgressed regions.

