## Additional File 1 for "Genomic evidence for recurrent genetic admixture during domestication mediterranean olive trees (*Olea europaea*)"

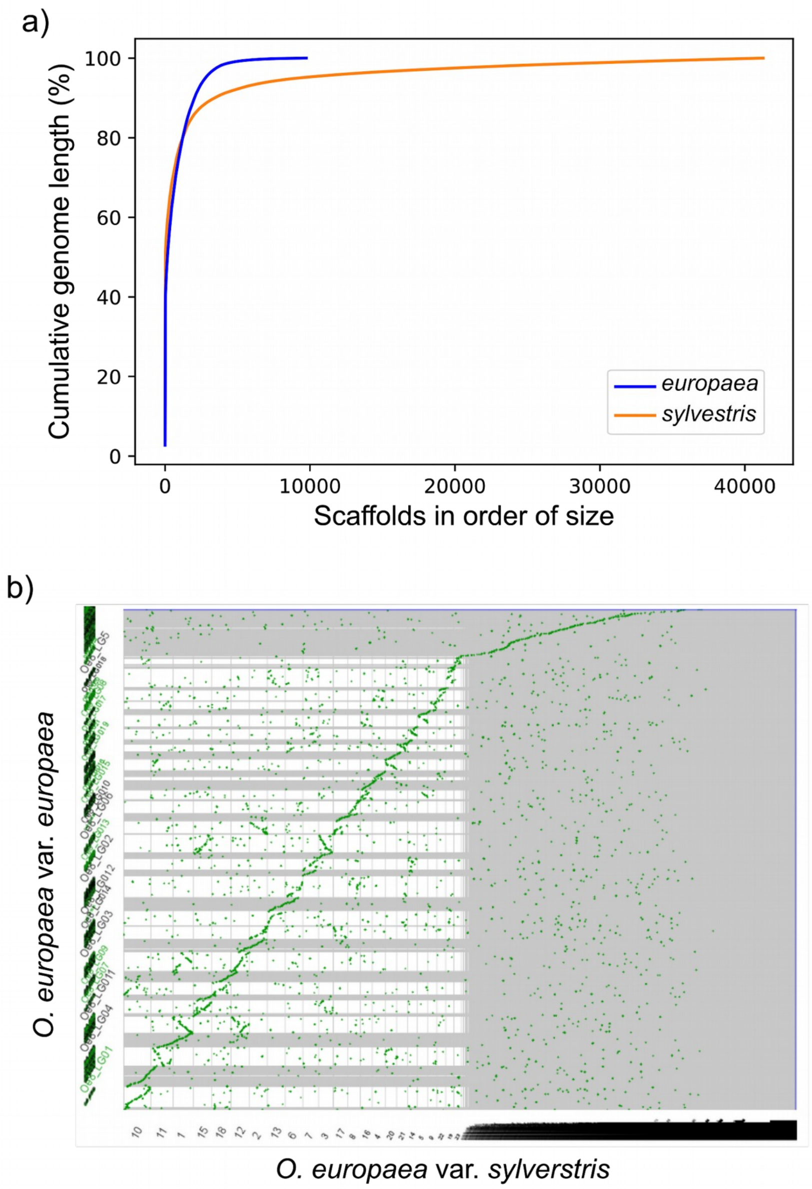


**Fig. S1. Genome comparison of *sylvestris* and *europaea*. (a)** Cumulative genome length per scaffold ranked in order of size for the genome assembly of *europaea* (red) and *sylvestris* (blue). A straight vertical line represents a perfect genome assembly. The horizontal plateaus indicates many small scaffolds. The top right end of each curve shows the total number of scaffolds. **(b)** Syntenic plot of the genome of *europaea* against *sylvestris* generated by SynMap.

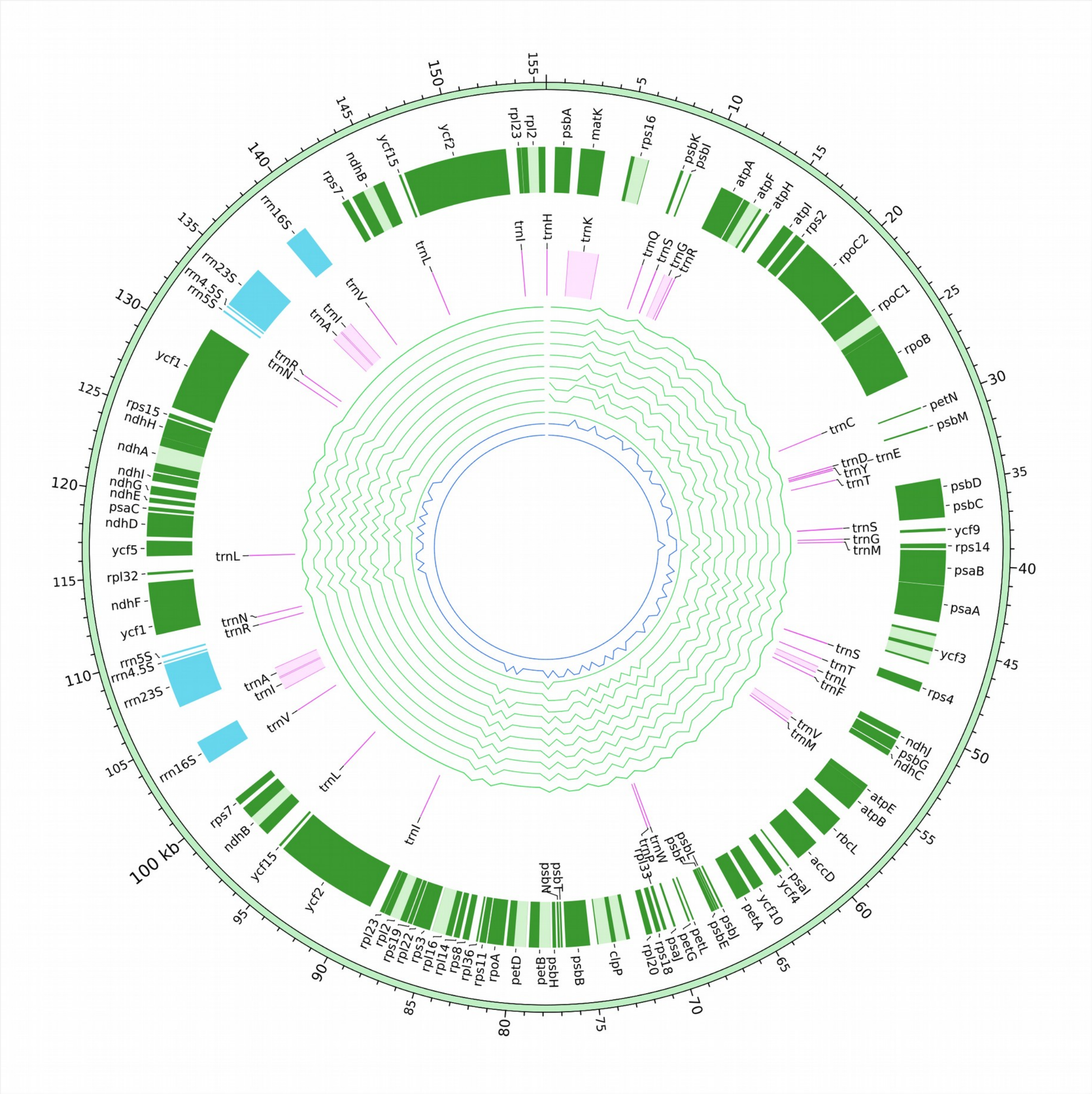


**Fig. S2. Plastid genome of the cultivar Farga.** Protein coding genes are shown in green, rRNAs in light blue, and tRNAs in purple. The SNPs are shown per each individual included in this study in the following order starting from outside: 'Arbequina', 'Picual', 'Beladi', 'Sorani', 'Koroneiki', 'Frantoio', 'Lechin de Granada', 'Lechin de Sevilla', ' Megaritiki', 'Chemlal de Kabilye', *sylvestris*-S, *sylvestris*-T, *maroccana*, *cerasiformis*, *guanchica*, *laperrinei*, *cuspidata*-R, *cuspidata*-S, *cuspidata*-I.

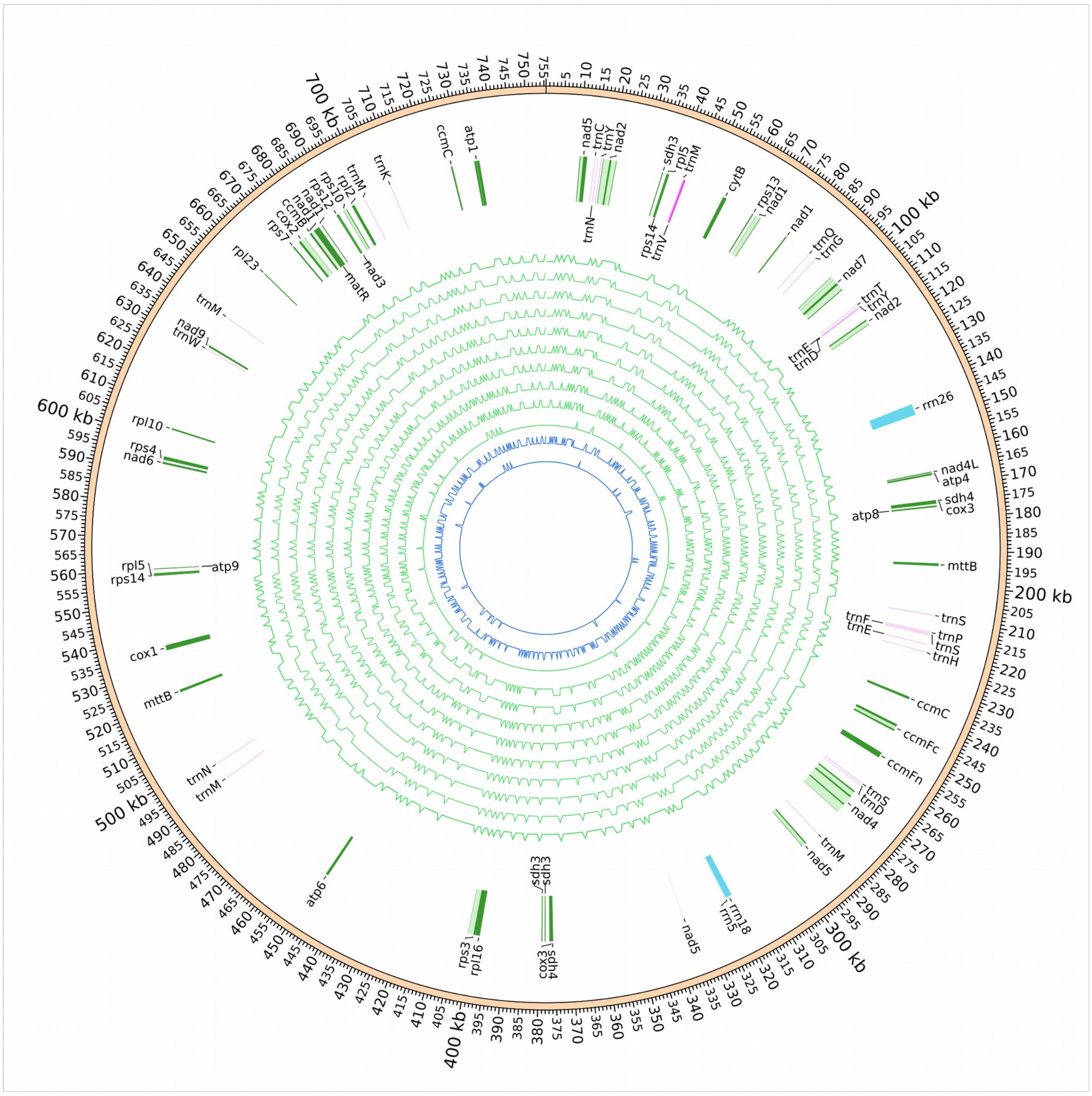

**Fig. S3. Mitochondrial genome of the cultivar Farga.** Protein coding genes are shown in green, rRNAs in light blue, and tRNAs in purple. The SNPs are shown per each individual included in this study in the following order starting from outside: ‘Arbequina’, ‘Picual’, ‘Beladi’, ‘Sorani’, ‘Koroneiki’, ‘Frantoio’, ‘Lechin de Granada’, ‘Lechin de Sevilla’, ‘ Megaritiki’, ‘Chemlal de Kabilye’, *sylvestris*-S, *sylvestris*-T, *maroccana*, *cerasiformis*, *guanchica*, *laperrinei*, *cuspidata*-R, *cuspidata*-S, *cuspidata*-I.

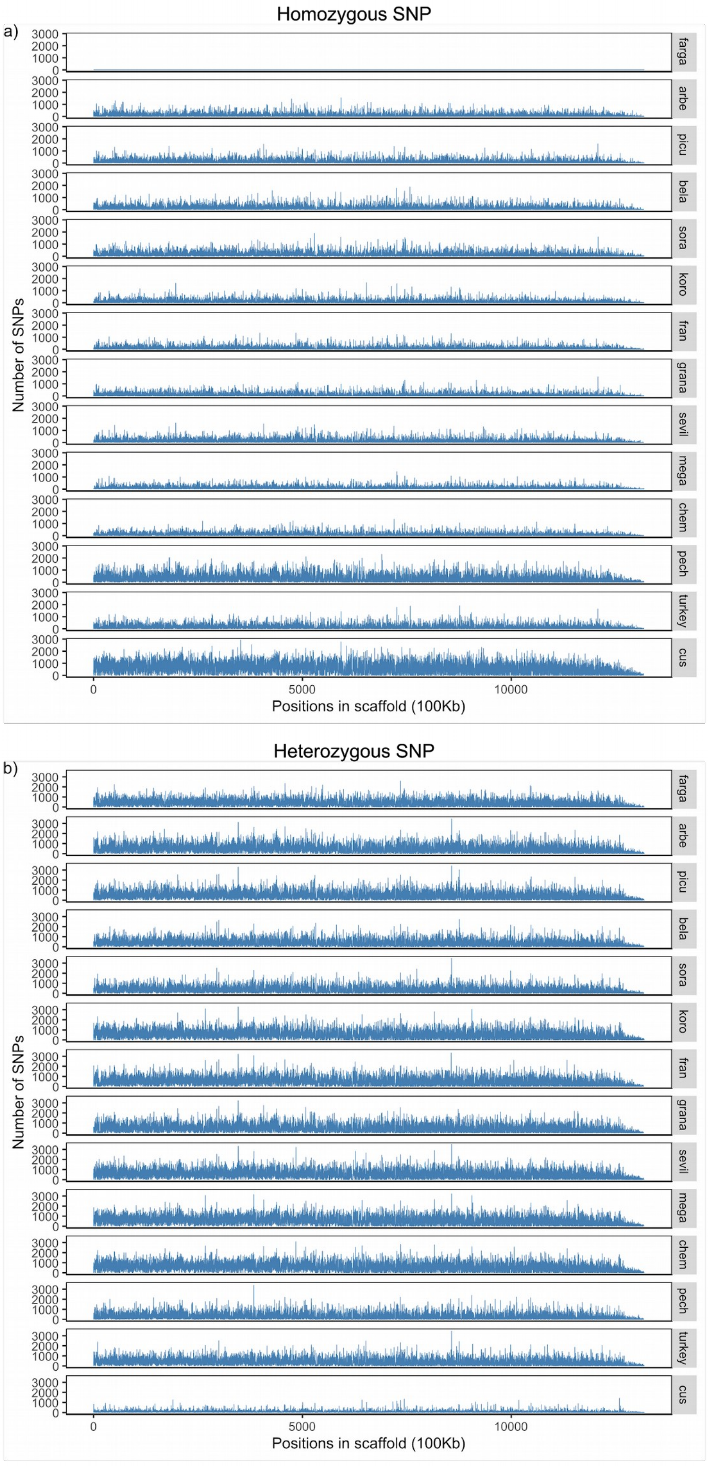


**Fig. S4. SNP distribution along the nuclear genome in windows of 100 Kb. (a)** homozygous SNPs, **(b)** heterozygous SNPs. Since cv. Farga was used as a reference genome, we do not expect homozygous SNPs for this sample.

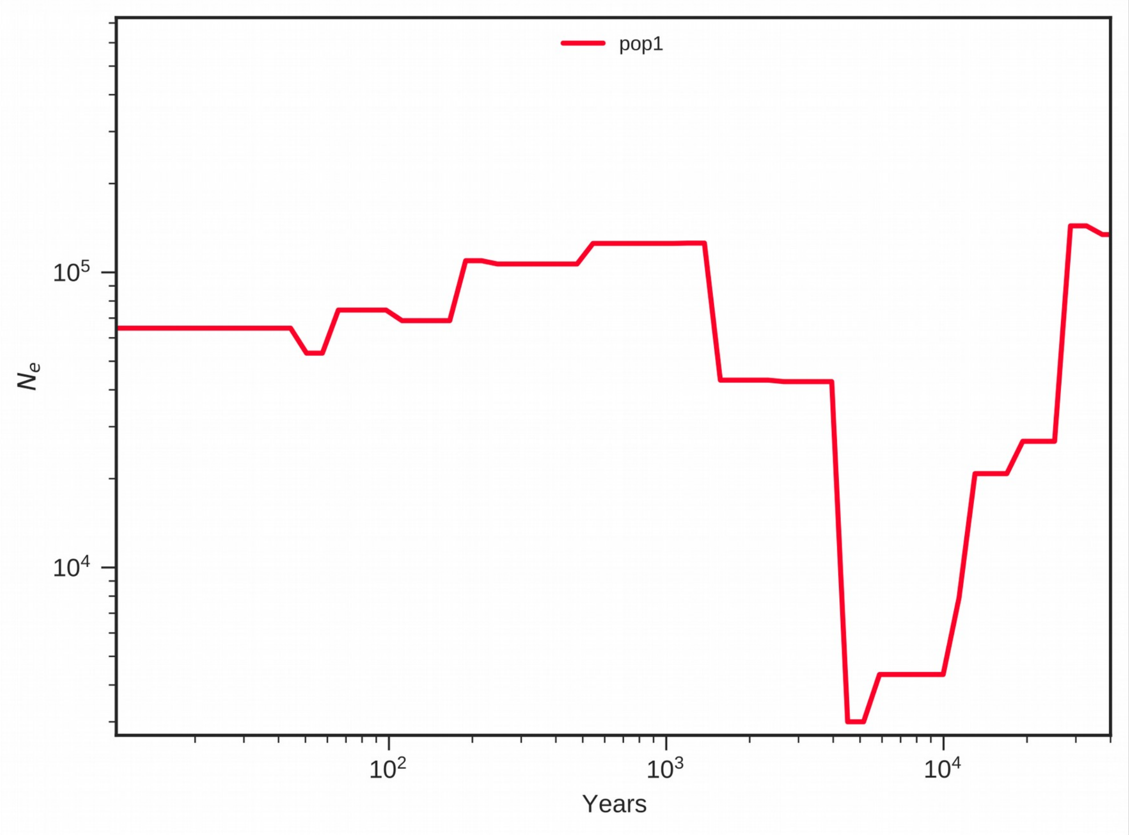


**Fig. S5. SMC++ results for inferring population size histories in clutivated olives.** A generation time of 20 years was used to convert coalescent scaling to calendar time.

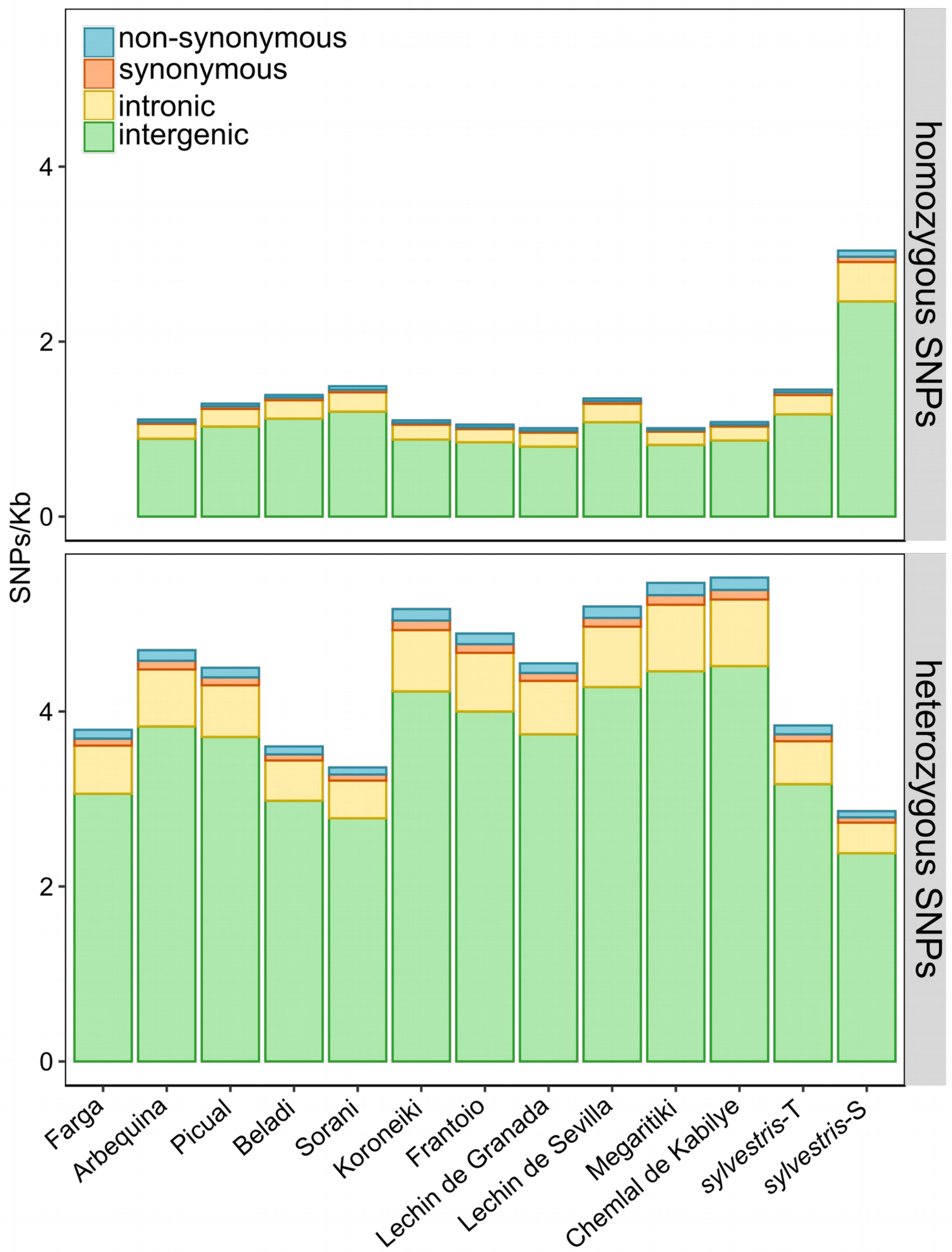


**Fig. S6. Number of homozygous and heterozygous SNPs (SNPs/Kb) in the intergenic, intronic and coding region of the genome.** The coding region was divided according to the changes that the allele can produce (synonymous and nonsynonymous).

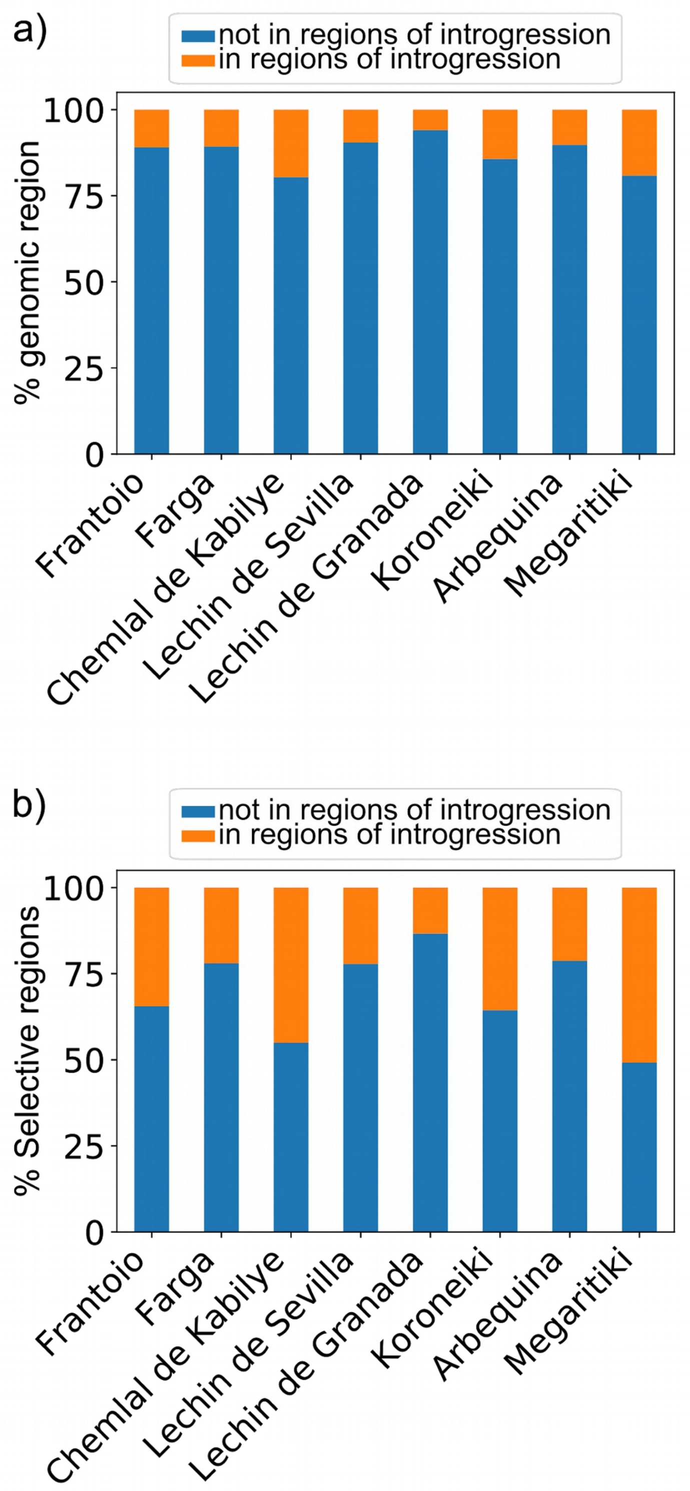


**Fig. S7. a)** Plot showing the percentage of genomic regions with introgression (D-statistic >0.5). **b)** Plot showing the percentage of selective sweeps that are present in introgressed regions.
